# *In vitro* and *in vivo* immunogenicity assessment of protein aggregate characteristics

**DOI:** 10.1101/2022.07.06.498969

**Authors:** Camilla Thorlaksen, Heidi S. Schultz, Simon K. Gammelgaard, Wim Jiskoot, Nikos S. Hatzakis, Flemming S. Nielsen, Helene Solberg, Vito Foderà, Christina Bartholdy, Minna Groenning

## Abstract

The immunogenicity risk of therapeutic protein aggregates has been extensively investigated over the past decades. While it is established that not all aggregates are equally immunogenic, the specific aggregate characteristics which are most likely to induce an immune response, remain ambiguous. The aim of this study was to perform comprehensive *in vitro* and *in vivo* immunogenicity assessment of human insulin aggregates varying in size, structure and chemical modifications, while keeping other morphological characteristics constant. We found that flexible aggregates with highly altered secondary structure were most immunogenic in all setups, while compact aggregates with native-like structure were found to be immunogenic primarily *in vivo*. Moreover, sub-visible (1-100 µm) aggregates were found to be more immunogenic than sub-micron (0.1-1 µm) aggregates, while chemical modifications (deamidation, ethylation and covalent dimers) were not found to have any measurable impact on immunogenicity. The findings highlight the importance of utilizing aggregates varying in few characteristics for assessment of immunogenicity risk of specific morphological features and provides a universal workflow for reliable particle analysis in biotherapeutics.

## Introduction

Biotherapeutics are an important and growing class of medicines, however they may lead to immune responses, such as anti-drug antibody (ADA) formation, which can compromise product efficacy and patient safety (Rosenberg & Sauna, 2018). Many patient- and product-specific factors are involved in the onset and progression of immunogenicity (Dingman & Balu-Iyer, 2019; Schellekens, 2005), and protein aggregation has been reported to be one of the major contributors to unwanted immunogenicity of biotherapeutics (Rosenberg, 2006). Examples of marketed products where ADA formation in patients were attributed to the presence of aggregates in the formulation are recombinant human erythropoietin (EPO) and interferons (Barnard *et al*, 2013; Kotarek *et al*, 2016; Seidl *et al*, 2012). Aggregation is an inherent capability of proteins and can occur at several stages during the product life-cycle, such as manufacturing, storage and patient handling (Gerhardt *et al*, 2014; Her & Carpenter, 2020; Randolph *et al*, 2015; Ueda *et al*, 2019; Walchli *et al*, 2020). Depending on external conditions, such as formulation (pH, ionic strength, etc.) and applied stress (shear force, temperature, primary packaging, etc.), the protein can undergo conformational changes leading to aggregates with various morphologies and physicochemical properties (Joubert *et al*, 2011; Mahler *et al*, 2009). However, it is currently not well understood how specific characteristics of protein aggregates impact immunogenicity (Bee *et al*, 2012; Rosenberg, 2006).

Current studies in the field have led to ambiguous results regarding which aggregate features contribute to drug immunogenicity (Hermeling *et al*, 2004). Some studies have suggested that highly aggregated samples are the most immunogenic (Barnard *et al*., 2013; Joubert *et al*, 2012), whereas others have found that even minute amounts of aggregate present in the therapeutic product seem to impact the risk of immunogenicity (Ahmadi *et al*, 2015). Another feature that has been widely discussed is stress-induced chemical modifications, such as oxidation. Some studies found oxidized species to be highly immunogenic (Boll *et al*, 2017; Filipe *et al*, 2012; van Beers *et al*, 2011), whereas others did not find evidence for this (Moussa *et al*, 2016a). The role of other chemical modifications, such as deamidation, is still not well described (Hermeling *et al*., 2004). Linking specific aggregate sizes to immunogenicity has likewise been evaluated. In this regard, all sizes ranging from sub-visible (1-100µm) to sub-micron particles (0.1-1µm) have been found to contribute to immunological reactions (Joubert *et al*., 2012; Kijanka *et al*, 2018; Telikepalli *et al*, 2015). Lastly, protein structure has been indicated as a possible factor impacting the immunogenicity. Misfolded protein aggregates might present different epitopes, which appear as foreign compared to native-like aggregates, with the risk of breaking self-tolerance. Nevertheless, both highly structurally-altered but also native-like aggregates have been shown to pose a risk (Freitag *et al*, 2015; Hermeling *et al*., 2004; Hermeling *et al*, 2006). The current studies mostly utilize monoclonal antibodies (mAbs) or pharmaceutical products already on the market, stressing them in various ways (heat, stirring, oxidation) and investigating the immunogenicity potential either *in vitro* or *in vivo*. However, the heterogeneity of the generated aggregate populations renders it difficult to disentangle the specific aggregate feature(s) impacting the immunogenicity potential (Boll *et al*., 2017). To get more clarity on this, a comprehensive and systematic approach is needed, where a detailed control of the aggregate formation and characteristics can help to directly couple the aggregate features to the immunological outputs.

Here we report on a systematic workflow to determine the profound immunogenic potential of specific aggregate characteristics. To achieve this we optimized protocols controlling the aggregation process and sample handling procedures, so that samples containing aggregates varying in few morphological features could be obtained (Thorlaksen *et al*, 2022a). We focused on four types of aggregates formed from recombinant human insulin. Two of the aggregates consisted of spherulites, which are micron-sized spherical superstructures, and consists of fibril-like protein material radially growing from a disordered core (De Luca *et al*, 2020; Krebs *et al*, 2004). The two spherulites had similar morphological appearance and size, but markedly varied in the β-sheet content (Thorlaksen *et al*, 2022b). The other two aggregates consisted of particulates, which are compact spherical structures ranging in size from nanometres to a few microns, where only minor secondary structural changes are induced upon aggregation (Foderà *et al*, 2014; Krebs *et al*, 2007). The particulates had similar morphological appearance, but markedly varied in size and also, to some extent, in secondary structure (Thorlaksen *et al*, 2022c). A detailed protocol for producing the tested aggregates as well as their extensive morphological characterization has been recently published by our group (Thorlaksen *et al*., 2022b; Thorlaksen *et al*., 2022c). Since highly relevant for the present publication, key characteristics are summarized in Table 1 together with new data quantifying the presence of chemical modifications. To assess the immunogenic potential of specific aggregate characteristics, the human DC response towards the aggregates was examined *in vitro* by measuring up-regulation of activation markers on the cell surface as well as inflammatory cytokine production. Activation of human CD4^+^ T-cells was evaluated based on cytokine secretion and cell proliferation upon co-culture with activated DCs pre-exposed to the aggregates. The aggregates were also tested *in vivo* by subcutaneous injections into BALB/c mice, where ADA formation towards human insulin was used as a direct measure of the immunogenicity. Our capacity to produce aggregate preparations with tuneable and controlled morphological features combined with comprehensive immunogenicity assessment *in vitro* and *in vivo* offered a detailed characterization of the link of aggregate features to immunological response. Despite the different criteria for immunogenicity between the *in vitro* and *in vivo* studies, we found a good correlation, where sub-visible aggregates with drastic structural alterations provided the highest immunological response in all cases.

**Table 1.**
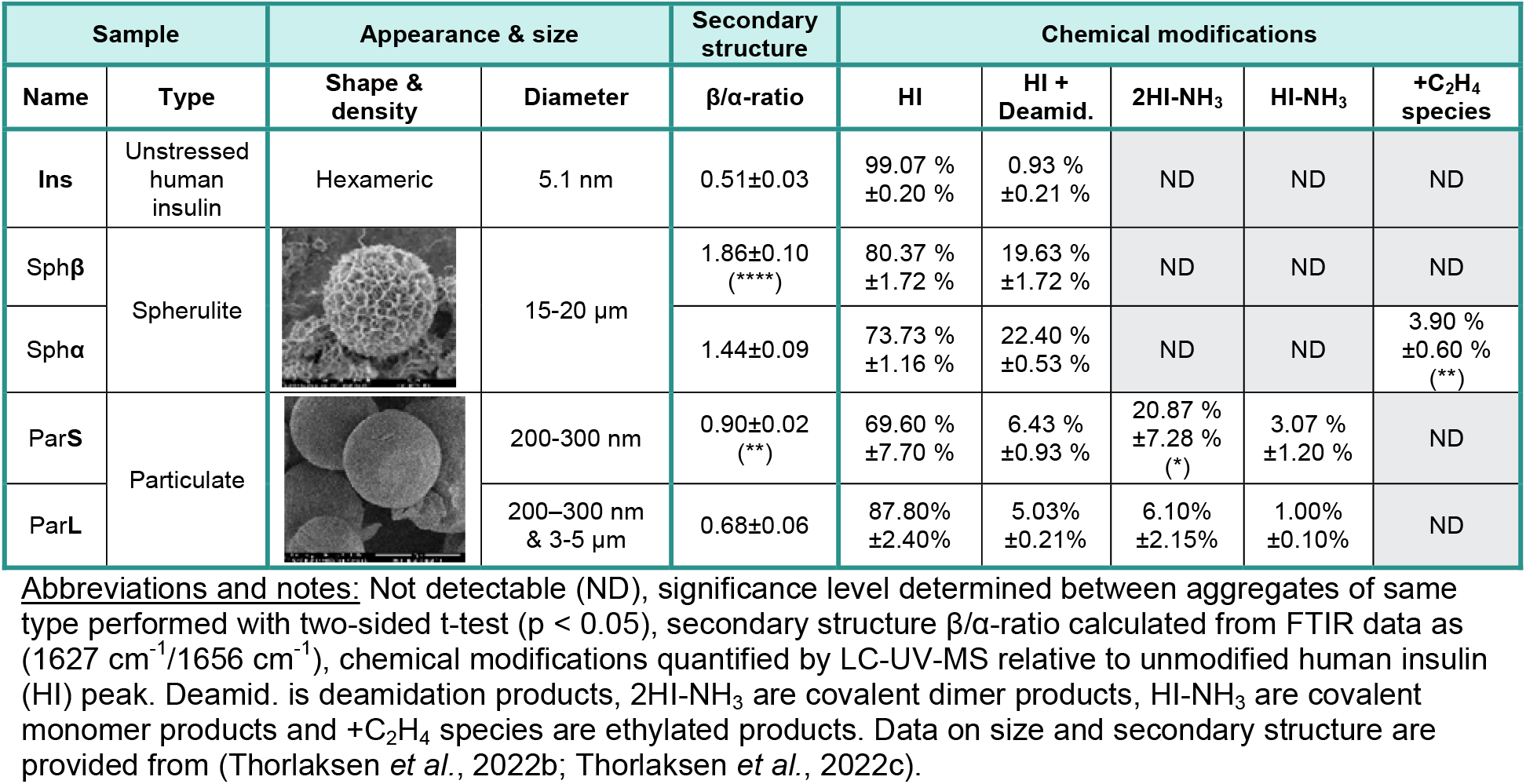
Overview of the tested samples and their characteristics.

| Sample |  | Appearance & size |  | Secondary structure | Chemical modifications |  |  |  |  |
| --- | --- | --- | --- | --- | --- | --- | --- | --- | --- |
| Name | Type | Shape & density | Diameter | $\beta/\alpha$-ratio | HI | HI + Deamid. | 2HI-NH<sub>3</sub> | HI-NH<sub>3</sub> | +C<sub>2</sub>H<sub>4</sub> species |
| Ins | Unstressed human insulin | Hexameric | 5.1 nm | 0.51±0.03 | 99.07 %<br>±0.20 % | 0.93 %<br>±0.21 % | ND | ND | ND |
| Sphβ | Spherulite | | 15-20 μm | 1.86±0.10<br>(****) | 80.37 %<br>±1.72 % | 19.63 %<br>±1.72 % | ND | ND | ND |
| Sphα | | | | 1.44±0.09 | 73.73 %<br>±1.16 % | 22.40 %<br>±0.53 % | ND | ND | 3.90 %<br>±0.60 %<br>(**) |
| ParS | Particulate |  | 200-300 nm | 0.90±0.02<br>(**) | 69.60 %<br>±7.70 % | 6.43 %<br>±0.93 % | 20.87 %<br>±7.28 %<br>(*) | 3.07 %<br>±1.20 % | ND |
| ParL | | | 200–300 nm & 3-5 μm | 0.68±0.06 | 87.80%<br>±2.40% | 5.03%<br>±0.21% | 6.10%<br>±2.15% | 1.00%<br>±0.10% | ND |

**Abbreviations and notes:** Not detectable (ND), significance level determined between aggregates of same type performed with two-sided t-test ( $p < 0.05$ ), secondary structure $\beta/\alpha$ -ratio calculated from FTIR data as ( $1627\text{ cm}^{-1}/1656\text{ cm}^{-1}$ ), chemical modifications quantified by LC-UV-MS relative to unmodified human insulin (HI) peak. Deamid. is deamidation products, 2HI-NH<sub>3</sub> are covalent dimer products, HI-NH<sub>3</sub> are covalent monomer products and +C<sub>2</sub>H<sub>4</sub> species are ethylated products. Data on size and secondary structure are provided from (Thorlaksen *et al.*, 2022b; Thorlaksen *et al.*, 2022c).

## Results

### Biophysical characterization of aggregates and unstressed insulin

Four aggregate samples were produced for immunogenicity assessment; two samples containing spherulites and two samples containing particulates. The two spherulite preparations were formed at acidic pH in either acetic acid or ethanol during heat stress, whereas the two particulate preparations were produced by varying the solution pH of 0.2 units, from pH 4.1 to 4.3 prior to heat stress. Before testing the aggregate samples in our immunogenicity assessment assays, they were comprehensively characterized and their original solvent was exchanged to phosphate buffer saline (PBS, pH 7.2-7.4) by dialysis. The complete characterisation findings are presented in two independent papers recently published by our group (Thorlaksen *et al*., 2022b; Thorlaksen *et al*., 2022c) and are summarized in Table 1 in terms of appearance, size and secondary structure.

The two spherulite preparations consisted of micron-sized (15-20 µm) spherical aggregates with a flexible structural appearance (open, loose and less compact structure), as visualized by the scanning electron microscopy (SEM) image in Table 1. Spherulites are known to consist of fibril-like material and are expected to exhibit a drastic increase in β-sheet formation compared to unstressed protein (Krebs *et al*., 2004). Interestingly, the β-sheet content can be decreased by forming the spherulites in ethanol (Sph**α**, β/α-ratio~1.44) compared to when formed in acetic acid (Sph**β**, β/α-ratio~1.86). The two particulate preparations consisted of compact spherical aggregates, as can be visualized by the SEM image in Table 1. The particulates formed at pH 4.1 (Par**S**) were in the nano size range (200-300 nm), while particulates formed at pH 4.3 (Par**L**) was a mixture consisting of both particles similar to the Par**S** sample and particulates in the micron range (3-5 µm). The nano-sized particulates in Par**S** had an increased β-sheet content (β/α-ratio~0.90), when compared to the micron-sized particulates found in Par**L** (β/α-ratio~0.68). Nevertheless, the particulates are considered more native in terms of secondary structure alterations (β/α-ratio < 1), than both the spherulites (see Fig. S1).

An unstressed human insulin sample (**Ins**) was used as a reference control in all the biological assays, to assess whether protein aggregation enhances the immunogenicity potential. The **Ins** sample was characterized by dynamic light scattering (DLS) to evaluate the protein oligomerization state in our formulation condition (1 mg/mL zinc-complexed human insulin in phosphate buffer saline, pH 7.2-7.4). An apparent hydrodynamic diameter of 5.1 nm indicates that the insulin is in its hexameric form, which is the expected dominant oligomerization state at physiological pH, when zinc is present (Schack *et al*, 2019). The secondary structure of the **Ins** sample was analysed with FTIR spectroscopy. Unstressed human insulin has a mainly α-helical structure (see Fig. S1). FTIR measurements on the **Ins** sample provided a calculated β/α-ratio of 0.51.

### Chemical characterization of aggregates and unstressed insulin

To form aggregates such as spherulites and particulates, the protein needs to be stressed under harsh conditions, such as low pH (below pH 7) and high temperature. These conditions could potentially induce chemical modifications on the human insulin molecules. We therefore investigated whether our aggregate preparations contained any chemically modified insulin molecules by dissociating the aggregates via a high pH treatment followed by a low pH treatment. We then analysed the samples (including **Ins** as reference) with reversed-phase liquid chromatography coupled to UV detection followed by mass spectrometry (LC-UV-MS).

Interestingly, we found that the pH treatment, which was used to dissociate the aggregates for LC-MS analysis, induced degradation of the intermolecular disulphide bonds between the A-chain (AC) and B-chain (BC) in the insulin structure (see Fig. S2). Alkaline pH can degrade disulphide bonds via β-elimination, which releases H_2_S (Florence, 1980; Wang *et al*, 2010) as well as introduces scrambling (Nagy, 2013; Trivedi *et al*, 2009). Dissolved H_2_S has been shown to convert disulphide bonds into trisulfide bonds in proteins (Gu *et al*, 2010; Hemmendorff *et al*, 2007). In the case of human insulin, the β-elimination in the interchain disulphide bonds combined with the scrambling into intrachain bonds could produce the free AC and BC species observed during the alkaline conditions. Moreover, we observed variations in the quantity of the degradation products between the aggregate types. This could suggest that the degradation induced by the pH treatment is dependent on the compactness of the structure and thus solvent accessibility. The samples, which were tested in the biological assays, did not undergo this pH treatment and most likely do not contain the mentioned degradation products. These products were therefore excluded from the further data analysis.

The major constituent in all the analysed samples was unmodified insulin (HI > 70%), while a fraction of the aggregates contained covalent alterations, primarily deamidation and dimerization (see Fig. 1 and Table 1). In both spherulite preparations, ~20% of the insulin had been deamidated, whereas in Sph**α** additionally ~4% were ethylated (+C_2_H_4_ species). The ethylation was found to be a result of the ethanol present in the formulation upon stress application. In the particulate preparations deamidated insulin (~5-6% for both preparations) was also found. Furthermore, two types of crosslinking products were found: the monomer either formed intramolecular crosslinking to preserve the monomeric status (HI-NH_3_) or two chemical groups from each monomer formed a covalent dimer through intermolecular crosslinking (2HI-NH_3_). Insulin dimers corresponding to 2HI-NH_3_ are common chemical degradation products that have previously been characterized (Darrington & Anderson, 1995; Hjorth *et al*, 2015). The crosslink sites were identified between the AC at Asn21 and BC at either the N-terminus or Lys29. In the two particulate preparations we observed a small amount of HI-NH_3_ products (1-3%), which were statistically indistinguishable. However, Par**S** was found to have a significantly higher amount of 2HI-NH_3_ products (~21%) compared to Par**L** (~6%). This could indicate that the covalent dimerization occurs primarily during the formation of the nanosized particulates.

**Figure 1.**
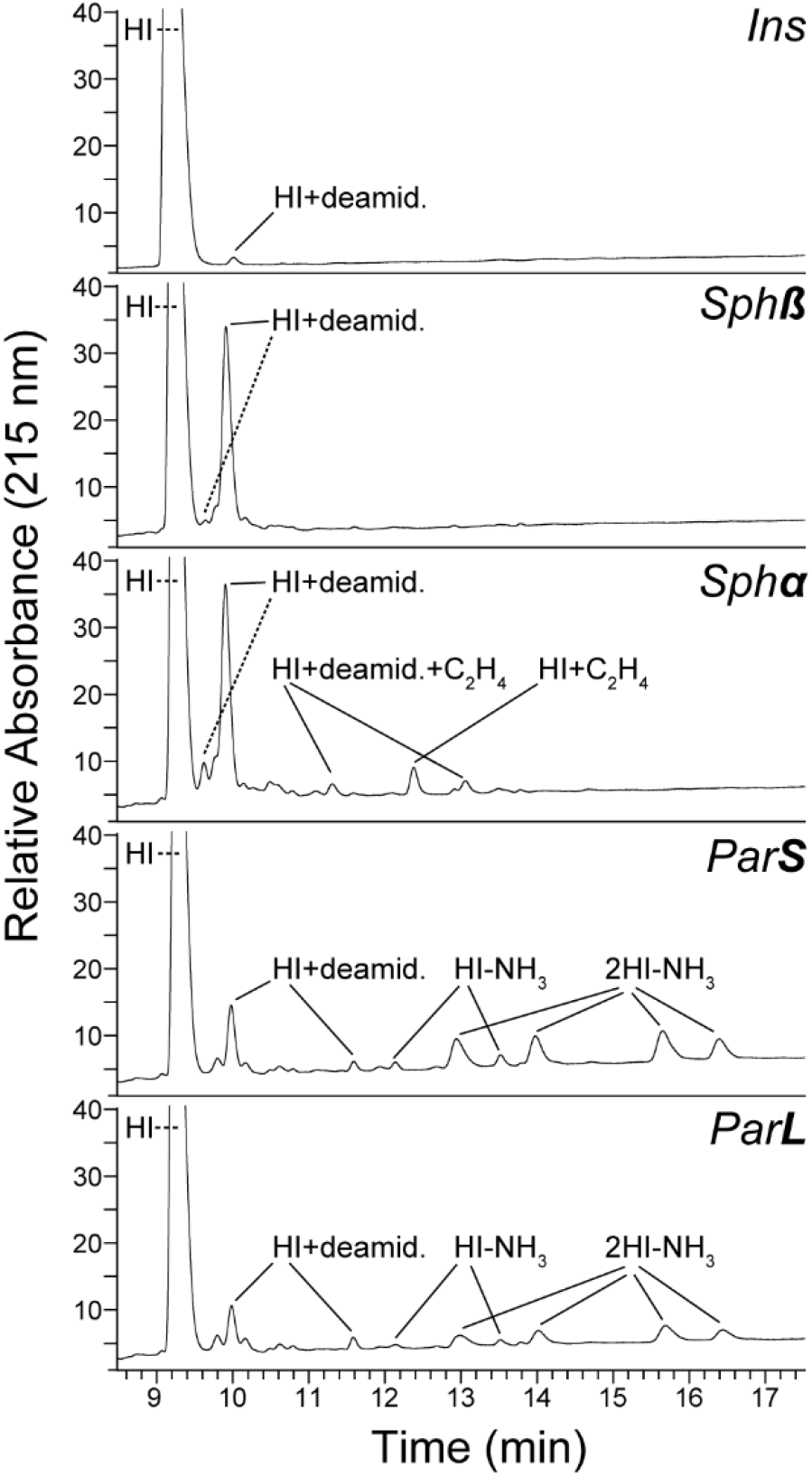
Identification of chemical modifications. Representative LC-UV-MS chromatograms of all tested compounds, where UV detection at 215 nm used for quantification of the chemical modifications formed during aggregation is illustrated. Chemical modification products observed after the main peak of unmodified human insulin (denoted HI) identified by mass spectrometry are annotated in the chromatograms for each sample. Aggregates were dissociated before sample analysis utilizing a high pH treatment followed by a low pH treatment.

### In vitro immunogenicity assessment of the aggregates

Generation of high affinity ADAs is complex and initially involves uptake, processing and activation of antigen-presenting cells (APC), such as dendritic cells (DCs), by the drug (Duke & Mitra-Kaushik, 2020). We developed an *in vitro* protocol for testing the immunogenicity potential of the aggregates, which allowed us to evaluate different stages of immune activation, spanning both the innate (DC activation) and adaptive (CD4^+^ T-cell activation) immune response (illustration in Fig. S4). We aimed at including donors in our study with specific HLA haplotypes (see Table S1), which were capable of binding insulin-derived peptides as assessed by *in silico* peptide–MHC II binding prediction (see Fig. S3). We initially investigated whether our aggregate preparations had the ability to activate monocyte-derived dendritic cells (moDC) in a similar manner as seen for other biotherapeutics in previously published literature (Ahmadi *et al*., 2015; Morgan *et al*, 2019). Activated moDCs are characterized by up-regulation of co-stimulatory molecules and maturation markers, secretion of inflammatory cytokines and chemokines, and lastly presentation of T-cell epitopes by MHC-II. MoDC activation was evaluated based on surface expression of the CD4^+^ T-cell co-stimulatory molecules CD80, CD86 and CD40 and maturation marker CD83. Stimulation with the positive controls (KLH and LPS) resulted in a significant up-regulation of all markers, compared to Ins (see Fig. S5). Treatment with the aggregate preparations mostly led to minor increases in the surface marker expressions. Only stimulation with Par**S** led to a significant increase in CD80 and CD86 expression, compared to Ins. MoDC’s stimulated with Sph**β** also showed trends of an increase in the average stimulation index (SI) for CD80, CD86 and CD40, however not statistically significant (see Fig. S5B). Secretion of inflammatory cytokines (IL-1β, IL-6 and TNF-α) and chemokine (IL-8) in the supernatant of the stimulated moDCs was likewise examined. In general, a significant increase in cytokines and chemokine was observed for the positive controls (KLH and LPS), when compared to Ins. Upon stimulation with the aggregate preparations, only Sph**β** showed an increase in IL-1β and IL-6 secretion, the latter being statistically significant (see Fig. S6).

T-cell activation and subsequent interaction with a cognate B-cell is an important step for ADA formation. The ability of the aggregate-stimulated moDCs to induce T-cell activation was evaluated by cell proliferation and secretion of the effector cytokine INF-γ. The positive control (KLH) and Sph**β** induced significant T-cell proliferation, when compared to Ins (see Fig. S7). The other aggregate preparations did not induce any proliferation. INF-γ secretion was measured at single cell level by fluorospot analysis (see Fig. S8A). KLH induced a high INF-γ response in all donors. For the aggregates, we observed that both Sph**β** and Par**L** induced a significant INF-γ response, compared to Ins (see Fig. S8B). These results suggest that the aggregate-stimulated moDCs can facilitate activation of CD4^+^ T-cells.

To provide a relative immunogenicity risk ranking of the aggregates, we calculated the percentage of responding donors and the magnitude of response for each of the four readouts (see Fig. 2). The frequency of responding donors and the corresponding magnitude (mean SI of the responding donors) was determined at single donor level. The criteria for a positive response were defined based on assay sensitivity and in accordance with other published papers (Joubert *et al*, 2016; Morgan *et al*., 2019; Schultz *et al*, 2017). The response to the aggregates was compared directly with the response to Ins. For the two DC activation assays, we found that > 33% of the tested donors responded to Sph**β** and Par**S**. Par**S** upregulated surface marker expression in 53% of the analysed donors with a magnitude of response at 2.1 relative to Ins, where Sph**β** provided the highest cytokine secretion response with a magnitude of response at 40, relative to Ins. Moreover, 27-29% of the donors responded to Par**L** and less than 13% of the donors responded to Sph**α**. In the T-cell proliferation assay we observed that donors only responded to Sph**β** (25%), whereas the INF-γ secretion assay allowed for a differentiation of the response to the aggregates. Here we found that 50% of the donors responded to Sph**β** and 25% responded to Par**L**, while only 17% of the donors responded to Par**S** and Sph**α**. Not all responding donors correlated across assays. However, based on the *in vitro* results, Sph**β** appears to have the highest immunogenicity potential across all four assays, followed by Par**L**, Par**S** and lastly Sph**α**.

**Figure 2.**
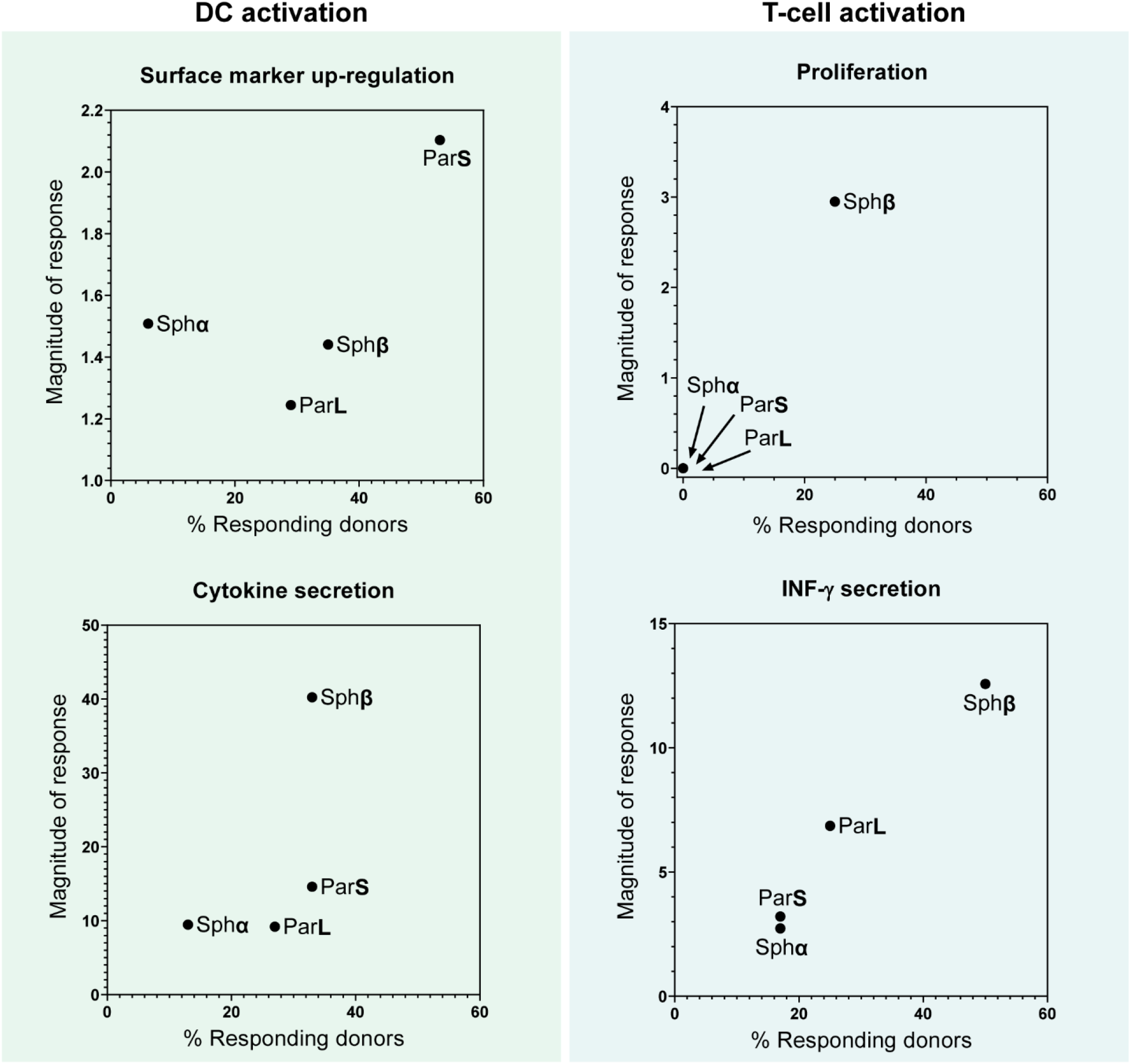
Percentage of responding donors vs. magnitude of response evaluated for all four *in vitro* immunogenicity assays. The % Responding donors are determined on single donor level for each assay and magnitude of response is the calculated average response (SI) for responding donors only. All aggregate samples are compared against the unstressed human insulin control (Ins).

### In vivo immunogenicity assessment of the aggregates

Even though human *in vitro* assays have become a valuable tool for assessing aggregate-related immunogenicity in terms of T cell activation, they do not address the complete and complex interplay between all components of the immune system, ultimately leading to activation of drug-specific B cells and production of high-affinity ADA responses (Jiskoot *et al*, 2016). Thus, BALB/c mice were repeatedly injected for 4 weeks with either Ins or aggregates to investigate whether the aggregates could enhance the development of anti-human insulin antibodies (see Fig. 3A). Although rodent *in vivo* models are generally considered poor predictors of human immunogenicity (Ratanji *et al*, 2014), we found it relevant to use them in this context as we were interested in examining adjuvant effects of the different aggregates rather than MHC-restricted responses to insulin-derived peptides. Human insulin is expected to be immunogenic in mice due to inadequate sequence homology to murine insulin and we therefore measured relative immunogenicity of the different aggregates rather than break of tolerance towards human insulin.

**Figure 3.**
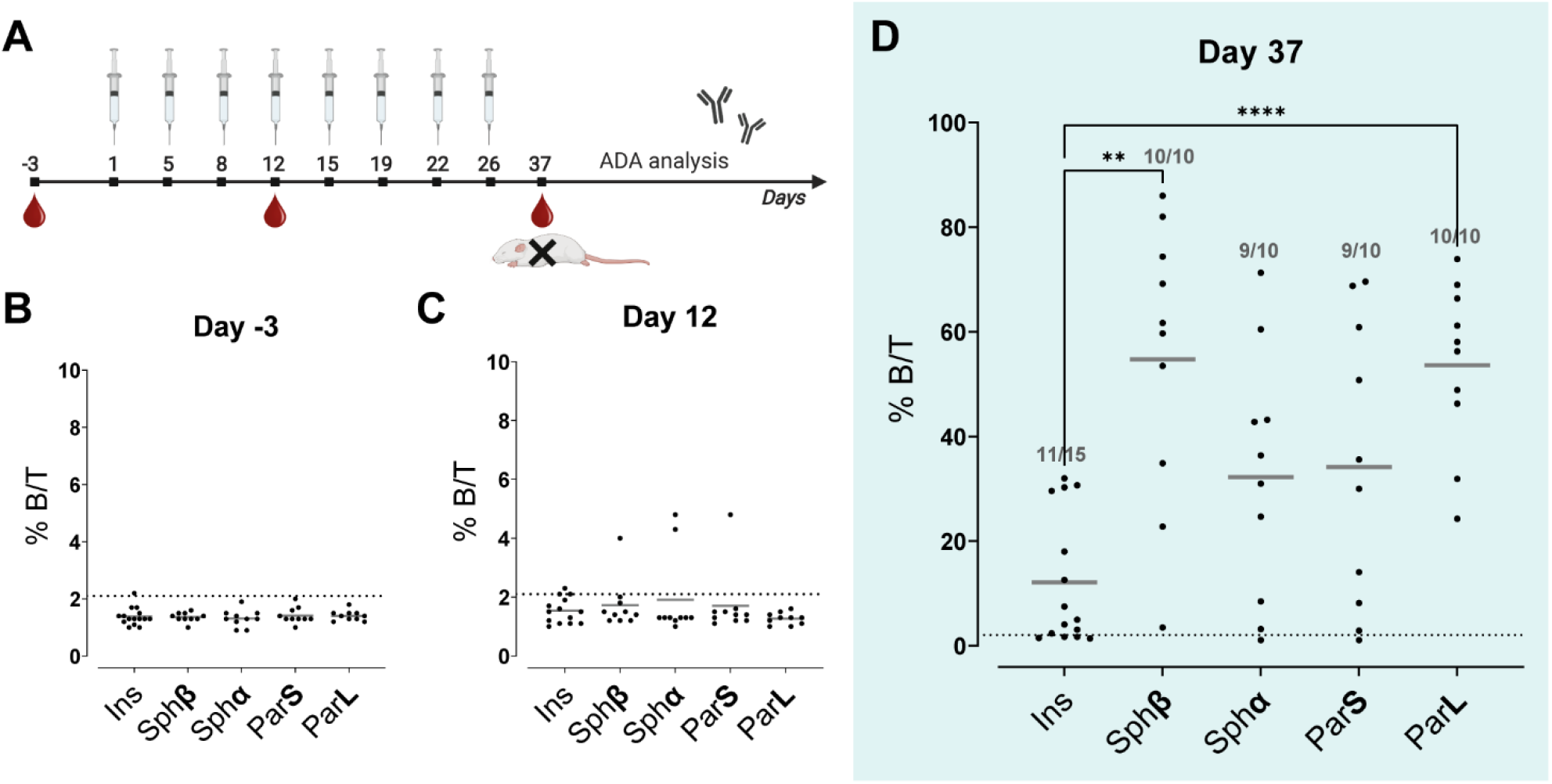
*In vivo* immunogenicity evaluation. A Graphical illustration of the experimental setup where BALB/c mice were injected SC twice-weekly with either Ins or aggregates. Blood plasma was collected at day −3, day 12 and day 37 and examined for the presence of ADAs by RadioImmuno Assay (RIA). The illustration was created with BioRender.com. B ADA assessment from blood plasma collected at day −3. C ADA assessment from blood plasma collected at day 12. D ADA assessment from blood plasma collected at day 37. Data information: The stippled line defines the “cut-off” point which is the average ADA response from day −3 plus 3 standard deviations. All responses above the “cut-off” point is considered positive. ADA titres are defined as %B/T in RIA. Significance level is evaluated by a one-way ANOVA Dunnet’s T3 multiple comparisons test.

The baseline levels of anti-human insulin antibodies in the mice were found to be similar to the assay blank controls and allowed calculating a “cut-off” point (%B/T = 2.1), from which we could define positive responses (see Fig. 3B). After 12 days of injections, no significant ADA responses were observed for either of the tested samples (see Fig. 3C). However, 11 days after the last injection, 49 out of 55 mice had developed anti-human insulin antibodies against our test samples (see Fig. 3D). Ins induced ADA responses in 11/15 mice correlating with the lack of full sequence homology between human and mouse insulin and consequently lack of central tolerance towards human insulin in mice. Sph**β** and Par**L** provided a statistically significant enhancement of the ADA response, compared to Ins both in terms of the frequency of responding mice and the magnitude of the response. Further analysis of the aggregate samples showed that the magnitude of the response induced by Par**L** was significantly higher than that induced by Sph**α** and Par**S**. The magnitude of response induced by Sph**β** was significantly higher than Sph**α**, but comparable to the response induced by Par**S**. Likewise, Sph**β** and Par**L** induced comparable magnitude of ADA responses. The results suggest that Sph**β** and Par**L** have the highest immunogenicity potential of the tested aggregates, followed by Par**S** and Sph**α**.

## Discussion

Aggregates are a major concern due to their potential to enhance the immunogenicity risk of a biotherapeutic protein. Yet the specific aggregate characteristics (e.g. size, secondary structure, chemical modifications) that are most likely to provoke an immune response are still ambiguous. To acquire this knowledge, we performed a comprehensive immunogenicity assessment of aggregates with well-defined features. This was possible as our earlier studies offered a detailed understanding and control of the production of human insulin aggregates with tuneable morphological characteristics. Even though healthy donors should be tolerant to human insulin, low amounts of insulin autoantibodies have been found in healthy donors indicating that some degree of break of tolerance can occur, which aggregates may accelerate (Williams *et al*, 1997). For our *in vitro* studies, we primarily included donors with specific human leukocyte antigen (HLA) haplotypes, HLA-DQ, suggested to play a role in the selection and activation of autoreactive T cells to increase the likelihood of provoking a T-cell response. By *in silico* peptide:MHC II binding prediction, we confirmed binding of insulin-derived peptides from the A and B chain to the selected HLA alleles, which is considered a pre-requisite for T-cell activation and subsequent ADA development.

DC activation is a decisive factor for whether a subsequent adaptive response, here measured as T-cell activation, is likely to be induced. Aggregates have the potential of providing an adjuvant effect by ensuring a larger delivery of epitopes for the T-cell due to the increased amount of protein molecules taken up and degraded by the DCs. Currently there is not much literature on the features protein aggregates should possess to obtain optimal DC uptake and presentation, or the activation mechanism. From vaccine research, parameters such as size, shape and rigidity (compact vs. flexible) are important for orchestrating an immune response (Benne *et al*, 2016). Some studies found that compact aggregates are more readily taken up by DCs than flexible particles (Christensen *et al*, 2012), and that larger particles (>500 nm) could enhance T-cell activation compared to smaller particles (Brewer *et al*, 2004). We would intuitively expect that the compact nanometre size particulates (200-300 nm) in the Par**S** and Par**L** preparations might be optimal for DC uptake, but not necessarily lead to a potent T-cell activation. In contrast, the spherulite preparations having a more flexible structure might activate the DCs via other routes, like receptors that recognize foreign repetitive motifs such as pathogen-associated molecular patterns (PAMPs) or by binding of complement components, which could facilitate subsequent T-cell activation (Moussa *et al*, 2016b).

We observed that Par**S** were able to up-regulate the expression of co-stimulatory surface markers in >50% of the tested donors, but only with a subtle magnitude of response. Moreover, Par**S** did not induce any T-cell activation, suggesting that this aggregate type is only slightly immunogenic. We observed no indication of DC activation for the Par**L** preparation. Nevertheless, Par**L** induced a potent adaptive response both *in vitro* (INF-γ secretion) and *in vivo* (ADA formation). The two particulate preparations have similar appearance, deamidation content and a native-like secondary structure (β/α-ratio<1), but Par**L** had a distinct micronsized particle population and Par**S** had a significantly higher content of covalent dimers (~21%). Par**L** induced an adaptive response unlike Par**S**, indicating that the presence of covalent dimers did not affect the immunogenicity potential. Thus our results suggests that the difference in immune activation is mainly due to the different size of these particles, where micron-sized particles (3-5 µm) provided a stronger immune activation than nanosized particles.

For the spherulite preparations, we found Sph**β** to be most immunogenic and Sph**α** to be the least immunogenic both *in vitro* (proliferation and INF-γ secretion) and *in vivo* (ADA titres). The two preparations have similar appearance, size and deamidation content, however Sph**α** contain less aggregated β-sheet compared to Sph**β** and was found to be slightly ethylated (~4%). Sph**α** did not lead to any measurable immune activation, hence we would not expect the ethylation nor deamidation to impact the immunogenicity potential. Therefore we propose that it is the difference in secondary structural features (β/α-ratio) from 1.44 (Sph**α**) to 1.86 (Sph**β**), that is the main driver of the increased immune activation observed. Interestingly, these findings also suggest that there might be a threshold for how much the secondary structure would have to be altered before the aggregates possess an increased immunogenic risk.

Even though *in vitro* assays have become a valuable tool for predicting aggregate-related immunogenicity, they do not cover all components of the complex immunological response leading to ADAs, e.g. T-cell dependent B-cell activation and antibody production. Moreover, the organized repetitive epitopes of aggregates might activate the immune system in a T-cell-independent pathway by direct B-cell receptor binding or through cross-linking of several B-cell receptors (Ratanji *et al*., 2014; Sauerborn *et al*, 2010; Wang *et al*, 2012). In our *in vivo* studies, we observed the micron-sized particulates in the Par**L** preparation, which are compact and have the most native-like structure among the aggregates, to provide an equally strong ADA response as the flexible Sph**β** particles with the most altered secondary structure. The Sph**β** particles could induce a potent T-cell response *in vitro*, suggesting that the immune activation might be mainly driven by a T-cell dependent pathway for this particle type. For Par**L** particles, the ADA response measured *in vivo* was highly potent, whereas a less potent T-cell activation was observed (only INF-γ secretion). This could indicate that the immune activation might be driven by a T-cell independent pathway for this particle type. Thus we speculate whether the compactness and native-like structure of the Par**L** particles present repetitive conformational epitopes with an optimal spacing and rigidity for B-cell receptor cross-linking (Moussa *et al*., 2016b; Sauerborn *et al*., 2010). Further characterization of the specific ADA isotypes induced by the individual aggregates would have to be performed to provide more solid knowledge as to the specific mode-of-action.

The approach presented here can be applied to assess immunogenicity risk of aggregates formed from various therapeutic proteins (i.e. mAbs). To do so, a comprehensive knowledge of the selected protein’s aggregation pathways must be obtained along with optimization of sample preparation to ensure sample homogeneity (same morphology). Our results show that this task proves challenging even with a known antigen model system such as insulin, and that the complexity increases once the aggregates are assessed in the biological assays. Moreover, the recombinant human insulin used in this study was expressed in yeast cells and it cannot be excluded that a human insulin molecule expressed in a different mammalian or bacterial cell line, can lead to a different glycosylation pattern, that would give a different immunogenicity profile. From this approach, it is thus not possible to relate the results directly to clinical immunogenicity. Yet, we can rank the aggregate characteristics relative to each other. Even though the criteria for immunogenicity assessment were different for our *in vitro* and *in vivo* studies, the resulted ranking of the tested aggregates according to their immunogenicity potential was well correlated and are presented in Fig. 4.

**Figure 4.**
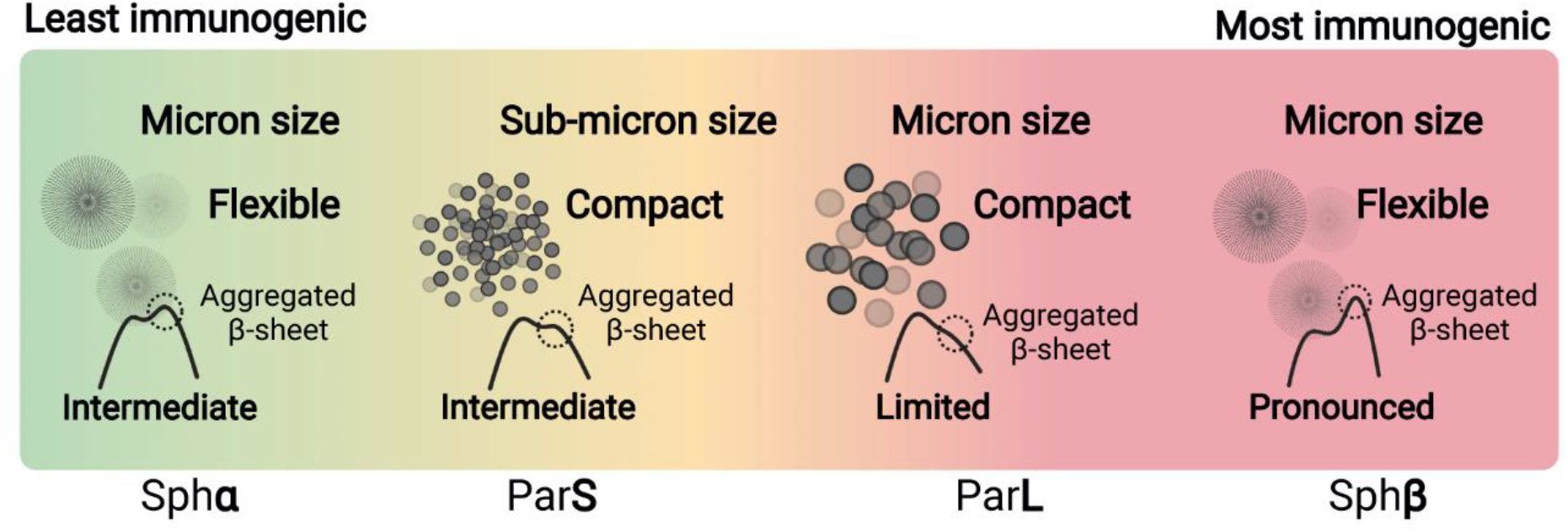
Ranking of immunogenicity potential of the tested aggregates. Graphical illustration of the ranking from least to most immunogenic (Sph**α** < Par**S** < Par**L** < Sph**β**) from our *in vitro* and *in vivo* immunogenicity assessment. The aggregates are presented by the morphological characteristics, which we found to impact the immunogenicity (size, appearance and secondary structure).

## Conclusion

Protein aggregation has been reported to be one of the major contributors to unwanted immunogenicity of biotherapeutics. We found that all of the tested aggregate populations were more immunogenic than unstressed human insulin and that the degree of immune activation was dependent on the specific aggregate features. Aggregates containing flexible micronsized particles with a highly altered secondary structure were found to be most immunogenic. Additionally, compact micron-sized particles with native-like structure were found to be immunogenic when examined *in vivo*. Particles in the sub-visible size range (1-100 µm) were found to be more immunogenic than particles in the sub-micron size range (0.1-1 µm). Lastly, we didn’t find any indications that chemical modifications (deamidation, conjugated dimers or ethylation) to the insulin molecule would impact the immunogenicity potential. We found a good correlation for immunogenicity assessment in the *in vitro* and *in vivo* studies. This is very interesting as the criteria for immunogenicity assessment are different. Our studies suggest that distinct aggregate types can activate different immunological pathways, collectively contributing to drug immunogenicity, which highlights the complexity of predicting clinical immunogenicity.

The insights we attained here were only possible when detailed biophysical studies of aggregate formation and characteristics were combined with *in vitro* and *in vivo* immunogenicity examinations. This allowed us to augment our understanding on how diverse feature impart diverse physiological responses. The findings have implications for formulation optimization of novel biotherapeutic products as well as for investigative purposes upon product recall. We anticipate that our universal workflow can form the blueprint for future studies on diverse therapeutic proteins for gathering comprehensive knowledge on the morphological characteristics affecting immunogenicity of biotherapeutics. This would most certainly lead to a better understanding of, e.g., which size, shape and structure of inherent or intrinsic protein particles in the drug product will pose the largest risk with regard to immunogenicity.

## Materials and methods

### Sample preparations and characterization

#### Unstressed human insulin

Zinc-complexed lyophilized recombinant human insulin (rh-HI), kindly provided by Novo Nordisk A/S, were dissolved in 200 µL of 0.1 M HCl (Merck KGaA, Darmstadt, Germany), followed by addition of DPBS (Dulbecco’s phosphate buffered saline, sterile, w/o calcium and magnesium, pH 7.4, #14190-094, gibco, Life Technologies Limited, Paisley, UK) for a final concentration of 1 mg/mL for cell studies or 93 µg/mL (80 nmol/kg) for animal studies. The pH of the solution was adjusted to pH 7.1-7.4 with 1 M NaOH (Merck KGaA, Darmstadt, Germany) and filtered with a silicone- and latex free syringe (#4050-X00V0, Henke Sass Wolf, Tuttlingen, Germany) through a 0.02 µm inorganic membrane filter (Alumina-based, Anotop 10, #6809-1002, Whatman, Germany) (**Ins** sample). For cell studies, the unstressed sample was prepared fresh on the day of stimulation and stored in a 5 mL NUNC cryotube vial (#379146, Thermo Fisher Scientific, Nunc A/S, Roskilde, Denmark) at 5 °C until use. For animal studies, the unstressed sample was prepared once weekly and stored at 5 °C in 5 mL vials (#60.558.001, 57×15,3mm, Sarstedt AG & Co., Nümbrecht, Germany). The sample was used for two rounds of injections and hereafter discarded.

#### Spherulite samples

Two spherulite samples were prepared according to the procedure reported in (Thorlaksen *et al*., 2022b). Briefly, rh-HI was prepared in either 20% acetic acid (v/v from acetic acid glacial, Merck KGaA, Darmstadt, Germany), 0.3 M NaCl (Merck KGaA, Darmstadt, Germany) at pH 2.0 with final protein concentration of 1.75 mg/mL (Sph**β** preparation) or in 40% ethanol (v/v from ethanol absolute, Merck KGaA, Darmstadt, Germany), 0.25 M NaCl at pH 1.8 with final protein concentration of 5 mg/mL (Sph**α** preparation). Sample preparations were filtered through a 0.1 µm inorganic membrane filter (Alumina-based, Anotop 25, #6809-2012, Whatman, Germany) using silicone- and latex-free syringes (#4050-X00V0, Henke Sass Wolf, Tuttlingen, Germany) directly into 15 mL Nunc conical centrifuge tubes (#339651, Thermo Fisher Scientific, NY, USA) (Thorlaksen *et al*., 2022a). Hereafter, aliquotes of 1 mL sample solution were transferred to 1.5 mL Eppendorf tubes (#0030 120.086, Eppendorf AG, Germany) and placed into a thermomixer (ThermoMixer C, Eppendorf AG, Hamburg, Germany) at 55 °C, 0 rpm for 8 h.

#### Particulate samples

The particulate samples were prepared according to the procedure reported in (Thorlaksen *et al*., 2022c). In short, 20 mM sodium acetate (Merck KGaA, Darmstadt, Germany) solution was added to rh-HI for a final concentration of 5 mg/mL. The solution was first titrated to pH 2.2 with 1 M HCl to dissolve the rh-HI, followed by titration with 1 M NaOH to reach a final pH of 4.1 (Par**S** preparation) or 4.3 (Par**L** preparation). The particulate samples were filtered in a similar manner as described above for the spherulites and aliquotes of 1 mL sample solution were transferred to 1.5 mL Eppendorf tubes. Samples were stressed in a thermomixer (thermomixer comfort, Eppendorf AG, Hamburg, Germany) at 80 °C for 2.5 h.

#### Solvent exchange by dialysis

Samples of spherulites and particulates were dialyzed against PBS (sterile, w/o calcium and magnesium, pH 7.2-7.4) before using them in cell- and animal studies, as described in (Thorlaksen *et al*., 2022b). Dialysis was executed at 4 °C to ensure particle integrity throughout the procedure. Particulate samples were dialyzed for 24 h, whereas spherulite samples were dialyzed for 48 h by using 100 kDa MWCO Float-A-Lyzer G2 dialysis devices (#G235059, Spectra/Por, Spectrum Laboratories Inc., CA, USA). Devices were cleaned prior to dialysis according to manufacturer recommendations. After sample recovery, the samples were stored at 5 °C for up to two weeks in 15 mL Nunc conical centrifuge tubes (#339651, Thermo Fisher Scientific, NY, USA) and then discarded.

#### Determination of protein concentration in aggregated samples

To obtain an estimate of the final protein concentration in each sample after dialysis, the aggregates would need to be dissociated. Dissociation was obtained by increasing pH to above pH 12 with 1 M NaOH for 10 minutes in a small sample aliquot (500 µL) of spherulites or particulates. Hereafter, the pH was decreased to below pH 8 with 1 M HCl. Protein concentration was determined for each sample by absorbance at 276 nm using a SoloVPE instrument (C Technologies Inc., NJ, USA). The final protein concentration was adjusted to account for dilution from the pH treatment procedure (high pH treatment followed by a low pH treatment).

#### Dynamic light scattering

To ensure hexamer formation in the **Ins** preparation, dynamic light scattering measurements were performed by using a DLS Dynapro II (Wyatt technology, CA, USA). **Ins** samples were always measured without further preparation. For analysis, 25 µL sample/well (in triplicates) was transferred into a 384 black microplate with transparent bottom (#ABA210100A, Aurora Microplate Products, MT, USA). Optical tape (#2239444, Bio-Rad Laboratories, USA) was used to cover the microplate top before centrifugation at 4000 rpm, 25 °C for 5 minutes to remove air bubbles from wells. Then, 10 µL/well silicone oil (#A12728, Alfa Aesar, ThermoFisher, Germany) was added on top of each sample. The instrument was left to equilibrate to 25 °C before measurements were started. 20 acquisitions of 2 seconds were used for each sample. The Dynamics software (version 7.9.1.4) automatically applied a cumulant fit to the data, which was used to estimate the sample hydrodynamic radii (nm) and polydispersity (%Pd).

#### Scanning electron microscopy

Scanning electron microscopy measurements were carried out in high vacuum mode, the samples were thus dried before analysis. The dried material was added onto a specimen stage covered with carbon tape (Agar Scientific Ltd, Essex, UK) and sputter coated with 0.2 nm gold using a Leica Coater ACE 200 (Leica Microsystems, Wetzlar, Germany). Imaging was performed using a FEI Quanta 3D FEG Scanning Electron Microscope (Thermo Fisher Scientific, Hillsboro, OR, USA) with an Everhart-Thornley Detector (ETD) and high-resolution dual beam. The acceleration voltage was 2.00 kV for acquisitions of particulate samples and 5.00 kV for spherulite samples.

#### Fourier-transform InfraRed spectroscopy

The **Ins** sample was prepared at a final protein concentration of 5 mg/mL for secondary structure analysis using a Tensor II Confocheck FTIR spectrophotometer with BioATR II module (Bruker Corporation, MA, USA). A background spectrum of PBS buffer (pH 7.2-7.4) was automatically subtracted from the protein spectra by the OPUS software (version 7.5.18) during measurements. Absorbance spectra were acquired in the range 4000 cm^−1^ - 900 cm^−1^ at a 4 cm^−1^ resolution, where each spectrum was an average of 512 scans. Between protein samples, the ATR module was cleaned with deionized water and 70% ethanol (v/v). The amide I peak (1600 cm^−1^ – 1710 cm^−1^) was used to evaluate structural features.

#### Fourier-transform InfraRed data interpretation

To quantitatively compare secondary structural features between the different samples (unstressed and the aggregates), FTIR spectroscopy and FTIR microscopy data from (Thorlaksen *et al*., 2022b; Thorlaksen *et al*., 2022c) were adapted such that a ratio between alpha-helix content (λ=1656 cm^−1^) and beta-sheet content (λ=1627 cm^−1^) could be estimated. All spectra were baseline corrected in range 2300 cm^−1^ −1800 cm^−1^ and normalized to 1656 cm^−1^. Subsequently, the β/α-ratio was calculated as (A1627 cm^−1^ / A1656 cm^−1^).

#### Liquid chromatography - mass spectrometry

An Acquity Classic UPLC (Waters, UK) was coupled to an Orbitrap Fusion Lumos with the H-ESI source (Thermo Scientific, CA, USA). The injection quantities were 1 ug HI. Reversed-phase chromatography was performed in an Acquity UPLC CSH C18 column at 1.0 × 150 mm, 1.7 μm (Waters, UK), with 0.1% formic acid in water as solution A (LS118-1, Optima, Thermo Scientific, USA) and 0.1% formic acid in acetonitrile (LS120-1, Optima, Thermo Scientific, USA) as solution B. The gradient was increasing from 20% to 32% solution B over 20 min with a flowrate of 100 μL/min and column temperature of 55°C. UV detection was at 215 nm. For the mass spectrometry settings, electrospray ionisation was at 3.2 kV, AGC target at 200k, and resolution at 120,000. The remaining settings were default. Xcalibur QualBrowser 4.2 (Thermo Scientific, USA) was used to analyse the LC-UV-MS data and quantify the UV-detected areas. The covalent products were quantified in triplicates from three separate batches of aggregate production.

#### Endotoxin analysis of aggregates

All sample preparations were tested for endotoxin levels by a LAL (Limulus Amebocyte Lysate) assay in-house at Novo Nordisk prior to being used in the biological assays. Endotoxin levels >1.00 EU/mL were found for all preparations.

### Immunogenicity assessment

#### *In silico* peptide:MHCII binding prediction

##### Prediction of human insulin-derived peptide:HLA binding affinities

The binding affinities of human insulin chain A and B to the most frequent European and North American HLA-DRB1, DQ and DP alleles was assessed using the affinity prediction algorithm NetMHCIIpan 3.0 (www.cbs.dtu.dk/services/NetMHCIIpan/), a software that predicts peptide–MHC-II binding affinities (Nielsen *et al*, 2008). An Epibar plot was generated to show the 9-mer core-binding peptide to the individual HLA class II alleles. The colours correspond to the binding affinity of the given allele. Two epitopes in both the A-chain and B-chain of human insulin were found to bind various HLA-DR/DP alleles, and especially specific HLA-DQ alleles of interest, indicating that they are capable of binding insulin (see Fig. S3).

#### *In vitro* immunogenicity

##### Preparation of PBMCs and cell isolation

Blood was donated for research use by HLA-typed healthy volunteers via the Danish Blood Bank (Copenhagen, Denmark) under informed consent, according to protocol H-D-2008-113 and approved by the Danish Scientific Ethical Committee Region Hovedstaden (Legislative Order no. 402 of May 28^th^ 2008). Donations were fully anonymous to Novo Nordisk A/S employees. For the study, the aim was to include donors with HLA-DQ2 (DQA1*0501/DQB*0201) and HLA-DQ8 (DQA1*0301/DQB1*0302), which are HLA haplotypes known to play a part in type 1 diabetes (T1D) susceptibility (Erlich *et al*, 2008; Van Autreve *et al*, 2004). The complete donor list can be found in Table S1. PBMCs were purified from whole blood by density centrifugation using Ficoll-Plaque Plus (GE Healthcare, Uppsala, Sweden) in Leucosep™ tubes (#227290, Greiner Bio-One GmbH, Germany), followed by lysis of the red blood cells with RBC lysis buffer (eBioscience, San Diego, CA, USA). The PBMCs were washed twice in DPBS buffer (Dulbecco’s phosphate buffered saline, sterile, w/o calcium and magnesium, pH 7.4, #14190-094, gibco, Life Technologies Limited, Paisley, UK) prior to monocyte and CD4^+^ T-cell isolation. First, monocytes were isolated by positive selection using a CD14^+^ microbeads human kit (#130-050-201, Miltenyi, Bergisch Gladbach, Germany). T-cells were isolated by negative selection from the CD14^−^ fraction using a CD4^+^ T-cell isolation kit, human (#130-096-533, Miltenyi, Bergisch Gladbach, Germany). Both isolation procedures were performed automatically by using an AutoMacs Pro instrument (Miltenyi Biotech) according to the manufacturer’s description. Cell purity in the isolated fractions was assessed by flow cytometry analysis and was found to be 93.9±2.8% for monocytes and 88.0±4.9% for CD4^+^ T-cells. All the CD4^+^ T-cells and a fraction of monocytes were frozen for later use with CryoStor^®^ CS10 (#C2874, Sigma-Aldrich, MO, USA), slow-frozen at −1 °C/minute (CoolCell^®^ LX, Corning) and stored at −80 °C. A graphical illustration of the full assay protocol can be viewed in Fig. S4.

##### DC activation assay

Monocytes were differentiated into immature dendritic cells (iDCs) by culturing in CellGenix^®^ GMP DC serum-free medium (CellGenix GmbH, Freiburg, Germany) supplemented with PenStrep (100 UI/mL), GM-CSF and IL-4 (40 ng/mL) for 5 days at 37°C, 5% CO_2_ in Nunc™ EasYFlask™ 75 cm^2^ (Thermo Fisher Scientific, Roskilde, Denmark). iDCs were transferred to a 96-well plate (Nunclon™ Delta surface, Thermo Fisher Scientific, Roskilde, Denmark) at 1.5×10^5^ cells/well. The cells were either left in media (unstressed) or stimulated with 300 nM Keyhole Limpet Hemocyanin (KLH, #SRP6195, Sigma-Aldrich, MO, USA), 1 µg/mL Lipopolysaccharide (LPS, #L2880, Sigma-Aldrich, MO, USA) or 100 µg/mL test compound (sample overview in Table 1) for 24 h at 37 °C, 5% CO_2_.

##### mDC:T-cell co-culture

iDCs were transferred to a 12-well cell culture plate (Corning Inc., ME, USA) at 0.6×10^6^ cells/well and were either left untreated or stimulated with 300 nM Keyhole Limpet Hemocyanin (KLH) or 100 µg/mL test compound (sample overview in Table 1). After 4 h incubation at 37 °C, 5% CO_2_, a maturation cocktail containing TNF-α and IL-1β (10 ng/mL) were added to the wells and incubation continued for another 40-42 h. CD4^+^ T-cells were quickly thawed in warm (37 °C) optimizer medium (OpTmizer™ CTS™ with T-cell expansion supplement, L-glutamine and PenStrep) and added to the mature DCs (mDCs) in a 1:10 ratio (6×10^6^ cells/well). The mDC:T-cell co-culture was incubated for 6 days at 37 °C, 5% CO_2_.

##### T-cell restimulation assay

A fluorospot 96-well plate from a Human INF-γ/IL-2 Fluorospot^PLUS^ kit (Mabtech AB, Sweden) was washed x3 with PBS and then blocked with optimizer media-10% FCS according to manufacturer description at room temperature until use. Monocytes were quickly thawed in warm (37 °C) DC media and 2×10^4^ cells/well were transferred to a fluorospot plate for cytokine secretion analysis and to a 96-well plate (Nunclon™ Delta surface, Thermo Fisher Scientific, Roskilde, Denmark) for proliferation analysis. The monocytes were stimulated with either media (untreated), 300 nM Keyhole Limpet Hemocyanin (KLH) or 100 µg/mL test compound (sample overview in Table 1) for 4 h at 37 °C, 5% CO_2_. Then, mDC:T-cell co-culture cells were added at 2×10^5^ cells/well. Additionally, anti-CD28 (#340975, BD Biosciences, CA, USA) 1 µL/well (0.1 µM) was added to the fluorospot plate and to the plate for proliferation analysis, 1 µM Click-iT^®^ EdU (#C10418, Invitrogen, Life Technologies Corporation, OR, USA) was added. Incubation were continued for another 40-42 h at 37 °C, 5% CO_2_.

##### Fluorospot analysis

Two days after re-stimulation, the fluorospot plate was washed x5 with DPBS, followed by incubation with anti-INF-γ (7-B6-1-BAM) detection antibody in DPBS-0.1% BSA for 2 h. The plate was washed and incubated with anti-BAM-490 fluorophore-conjugate in DPBS-0.1% BSA for 1 h, while protected from light. Lastly, the plate was washed and spots were developed using fluorescence enhancer for 15 minutes. Dried plates were analysed with an Immunospot S6 Universal Analyzer (Cellular Technology Limited, OH, USA) using the build-in software.

##### Cytokine measurements

Plates from the DC activation assay were centrifuged at 1200 rpm, 4 °C for 2 minutes. The supernatant (100 µL/well) was transferred to a U-bottom well plate (#650201, Greiner Bio-One, USA) and stored at −80 °C until use. Multiplex analysis of cytokines (IL-1β, IL-6, IL-8 and TNF-α) on the supernatants was performed using a 10-spot MSD U-PLEX platform, biomarker group 1 human 4-assays (Meso Scale Discovery, MD, USA) according to the manufacturer instructions. In brief, U-plex linkers were mixed with their corresponding biotinylated antibodies in separate tubes and left for 30 minutes incubation. Stop solution was added, the tubes were vortexed and left for another 30 minutes incubation. The U-plex linker-coupled antibody solutions were mixed, transferred to the MSD plate and incubated for 1 hour with shaking. The MSD plate was washed x3 with DPBS-0.05% Tween 20, followed by sample addition (DC supernatants) and incubation for 1 h with shaking. Plate was washed x3 with DPBS-0.05% Tween 20 and a Sulfo-tag™ conjugated detection antibody solution, containing detection antibodies corresponding to each analyte, was added to the wells. The solution was left for 1 h incubation with shaking and washed x3 with DPBS-0.05% Tween 20. Lastly, read buffer was added and the plate was analysed on a MSD instrument (Meso Scale Discovery, MD, USA) by using the MSD Discovery Workbench software. Analyte concentration (pg/ml) was extrapolated from luminescence, based on a standard curve generated from multi-analyte calibrator standards. Data values that exceeded the limit-of-detection (lower- or upper-LOD) of the standard curve, was replaced with the calculated LOD for each analyte.

##### Flow cytometry

DC surface marker expression from the DC activation assay was assessed by multicolour flow cytometry using a panel of mouse anti-human monoclonal antibodies: FITC conjugated anti-CD80, APC conjugated anti-CD83, PE conjugated anti-CD86, V450 conjugated anti-CD40 (all from BD Biosciences, CA, USA) and APC-Cy7 conjugated LIVE/DEAD™ Fixable Near-IR Dead Cell Stain (#L10119, Invitrogen, Thermo Fischer Scientific, OR, USA). Negative controls were isotype matched control antibodies. Cells were transferred to a U-bottom well plate (#650201, Greiner Bio-One, USA), washed twice with DPBS-1% FCS and once with DPBS-10% FCS. To prevent unspecific antibody binding, the cells were first incubated for 10 min with human Fc block (#564220, BD Biosciences, CA, USA) in DPBS-10% FCS. The cells were stained with the antibody panel for 20 minutes and washed x3 with DPBS-1% FCS prior to flow cytometry analysis. For T-cell proliferation analysis, the cells were first extracellularly stained with a mouse anti-human monoclonal antibody APC-Cy7 conjugated LIVE/DEAD™ Fixable Near-IR Dead Cell Stain (#L10119, Invitrogen, Thermo Fischer Scientific, OR, USA), following a similar preparation procedure as just described for the DCs. Hereafter, the cells were click labelled intracellularly using the Click-iT^®^ EdU flow cytometry assay kit (#C10418, Invitrogen, Life Technologies Corporation, OR, USA) according to manufacturer instructions. Briefly, the cells were fixed and permeabilized by incubation for 15 minutes with Click-iT^®^ fixative. The plate was washed twice in Click-iT^®^ wash reagent and Click-iT^®^ reaction cocktail (Pacific Blue™ azide, CuSO_4_ and reaction buffer additive) were added to each well, followed by 30 minutes incubation. Lastly, the cells were washed in wash reagent before flow cytometry analysis. For compensation, the individual antibodies used for staining were mixed with one drop of negative control beads and anti-mouse IgG beads (BD Biosciences, CA, USA). For the dead cell stain, compensation was performed on negative control beads and anti-Rat/Hamster IgG beads (#552845, BD Biosciences, CA, USA). All flow cytometry assays were performed on a LSR FACSFortessa instrument (BD Biosciences, CA, USA), while Kaluza software (v.2.1) were used for data analysis.

##### Statistical analysis

Results from all four assays were plotted as ‘stimulation index’ (SI) vs. stimulation condition. The SI is calculated as: Response_condition_ / Response_untreated cells_. The significance level (p<0.05) was determined by a one-tailed t-test, assuming normality and unequal variance. To plot % Responding Donors vs. Magnitude of Response, a positive response was defined at single donor level in each assay. For DC surface marker data, a positive response was obtained if the SI was significantly up-regulated and p<0.05 in a one-tailed t-test for either one of the markers (CD80, CD83 or CD86). For multiplex cytokine data, a positive response was obtained if SI > 2 for all four cytokines. For fluorospot and proliferation analysis, a response was defined as positive if SI > 2 and p<0.05 by a one-tailed t-test. For all assays, KLH and LPS (if present) response were compared to untreated cells, whereas aggregates were compared to cells stimulated with unstressed insulin (Ins sample). The Magnitude of Response in all assays were calculated as the average SI for responding donors.

#### *In vivo* immunogenicity

##### Animal husbandry

The care and use of mice in these studies were conducted according to national regulations in Denmark and with experimental licenses granted by the Danish Ministry of Justice. The mice were housed under 12:12 light-dark cycle in humidity- and temperature controlled rooms with free access to standard chow (catalogue 1324, Altromin, Brogaarden, Denmark) and water throughout both the acclimation (12 days) and study period (37 days).

##### Animal study

A total number of 55 BALB/c mice (BALB/cAnNCrl, 7 weeks of age) purchased from Charles River Laboratories (Sulzfeld, Germany) were divided into 5 groups; A control group treated with unstressed human insulin (N=15) and 4 groups treated with one of the different aggregate types (N=10/group). The mice were injected SC twice weekly with 80 nmol/kg test compound at dose 5 mL/kg for 4 consecutive weeks using Injekt-F Luer Duo silicone- and latex-free syringes and 25Gx5/8” needles (#9166033V, B. Braun, Melsungen, Germany). The **Ins** sample was prepared weekly, whereas aggregate samples were prepared every 2 weeks. Samples were stored at 5 °C in 5 mL vials (#60.558.001, 57×15,3mm, Sarstedt AG & Co., Nümbrecht, Germany), but were equilibrated to room temperature before injections. The study design was inspired by a similar previously published study (Kijanka *et al*., 2018). Three days before the first injection (day −3), 200 µL blood was collected sublingual to obtain pre-dose samples for cut-point calculation of the basal ADA level. Midway through injections (day 12), another 200 µL blood was collected sublingual. The mice were sacrificed one week after the last injection (day 37). Here 500 µL blood was collected sublingual. To isolate plasma for ADA analysis, blood was sampled directly in tubes containing EDTA and put into ice water before centrifugation for 5 minutes, 6000 x g at room temperature. The stabilized plasma was kept in 0.7 mL Micronic tubes (Micronic, Lelystad, Netherlands) at −20 °C until use.

##### Anti-insulin antibody detection

The anti-drug antibody (ADA) responses were evaluated with Radioimmunoassay (RIA) adapted to detect anti-human insulin antibodies in mouse plasma. First, plasma samples were mixed with human insulin (rh-HI) tracer [Tyr^A14^(^125^-I] and assay buffer (0.04 M phosphate, 0.15 M NaCl, 0.5% w/v BSA, 0.25% γ-globulin, 0.01 M EDTA, pH 7.4) into Nunc mini Sorp tubes (Nunc A/S, Roskilde, Denmark) in duplicates. The mixture was vortexed for 10 seconds, followed by 24 h incubation at 5 °C to allow anti-insulin antibodies in the plasma to bind the rh-HI tracer. Second, 14.4% (w/v) PEG-6000 (#807491, Merck) in TBS buffer (0.01 M Trizma base, 0.15 M NaCl, 0.1% Tween 20, pH 8.6) was added to each tube to precipitate antibodies in the sample. The tubes were vortexed for 10-20 seconds and then centrifuged 30 minutes, 3000 rpm at 4 °C. The supernatant was discarded, and the precipitate was washed once with PEG-6000 in TBS followed by centrifugation for 15 minutes, 3000 rpm at 4 °C. Lastly, the supernatant was discarded before being placed into the gamma counter (Wizard2, 10 detector, PerkinElmer). Each tube was counted for 120 seconds. The measured cpm (count-per-minute) is proportional to the amount of bound anti-insulin antibodies bound to the rh-HI tracer in the sample. Counts were normalized to tubes with unprecipitated rh-HI tracer, called “total”. The results are given as bound(cpm) / total(cpm) * 100 (%B/T). A response was classified as positive if the %B/T value was equal or higher than the cut-point, defined as the upper 99.7^th^ percentile (3 SD) of the calculated %B/T for samples collected at day −3.

##### Statistical analysis

To statistically evaluate the potential differences in anti-insulin antibody response between unstressed human insulin and the aggregates, a One-Way ANOVA with Dunnett T3 (n < 50/group) multiple comparisons test assuming unequal SDs were applied to the data using GraphPad Prism software (v. 9.0.1).

## Acknowledgements

Authors acknowledge Novo Nordisk A/S for project funding and providing material for formation of protein aggregates, human cell studies and animal studies. VF and CT acknowledge VILLUM FONDEN for supporting the project via the Villum Young Investigator Grant “Protein Superstructures as Smart Biomaterials (ProSmart)” 2018−2023 (Grant 19175). NSH and CT acknowledge VILLUM FONDEN (Grant 18333) and NSH acknowledge Novo Nordisk foundation grants NNF16OC0021948 and NNF14CC0001. For use of SEM, authors acknowledge the Core Facility for Integrated Microscopy, Faculty of Health and Medical Sciences, University of Copenhagen.

## Author contributions

CT and HSS developed protocols, performed experiments and wrote the manuscript draft. SKG performed LC-UV-MS experiments. WJ and NSH supported with data interpretation and interesting scientific discussions. FSN planned and executed the animal studies. HS planned, executed and interpreted ADA analysis experiments. VF, MG and CB designed the research project and were responsible for the overall planning and management. All authors agreed on the data interpretation and contributed to the manuscript.

## Figure legends

**Figure S1.**
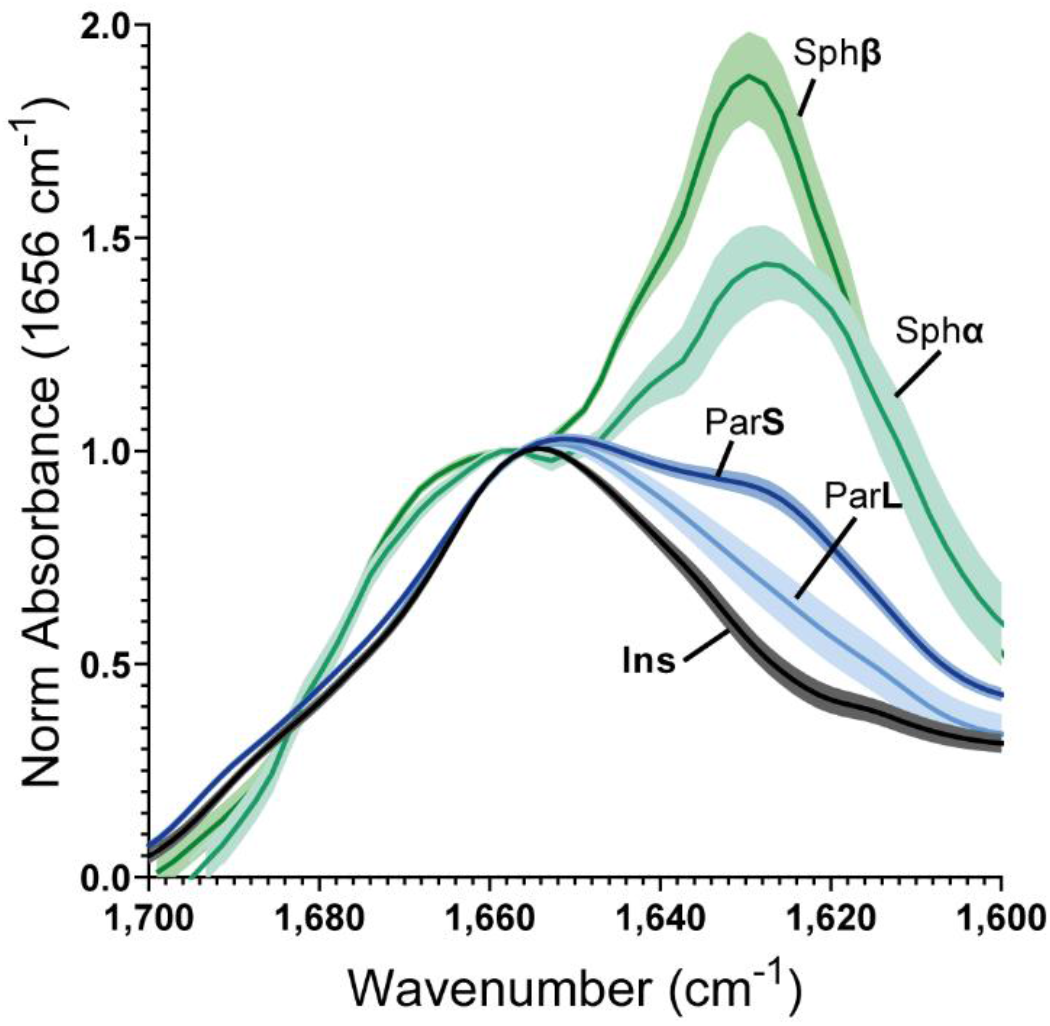
Secondary structure characterization. Fourier-Transform InfraRed spectra of **Ins** (black), Par**S** (dark blue), Par**L** (light blue), Sph**β** (dark green), Sph**α** (light green) normalized at 1656 cm^−1^ (α-helix content) to illustrate the difference in β-sheet content at 1627 cm^−1^ between the aggregate samples compared to the unstressed HI sample. Data for the aggregate populations has been adapted from previously published work (Thorlaksen *et al*., 2022b; Thorlaksen *et al*., 2022c).

**Figure S2.**
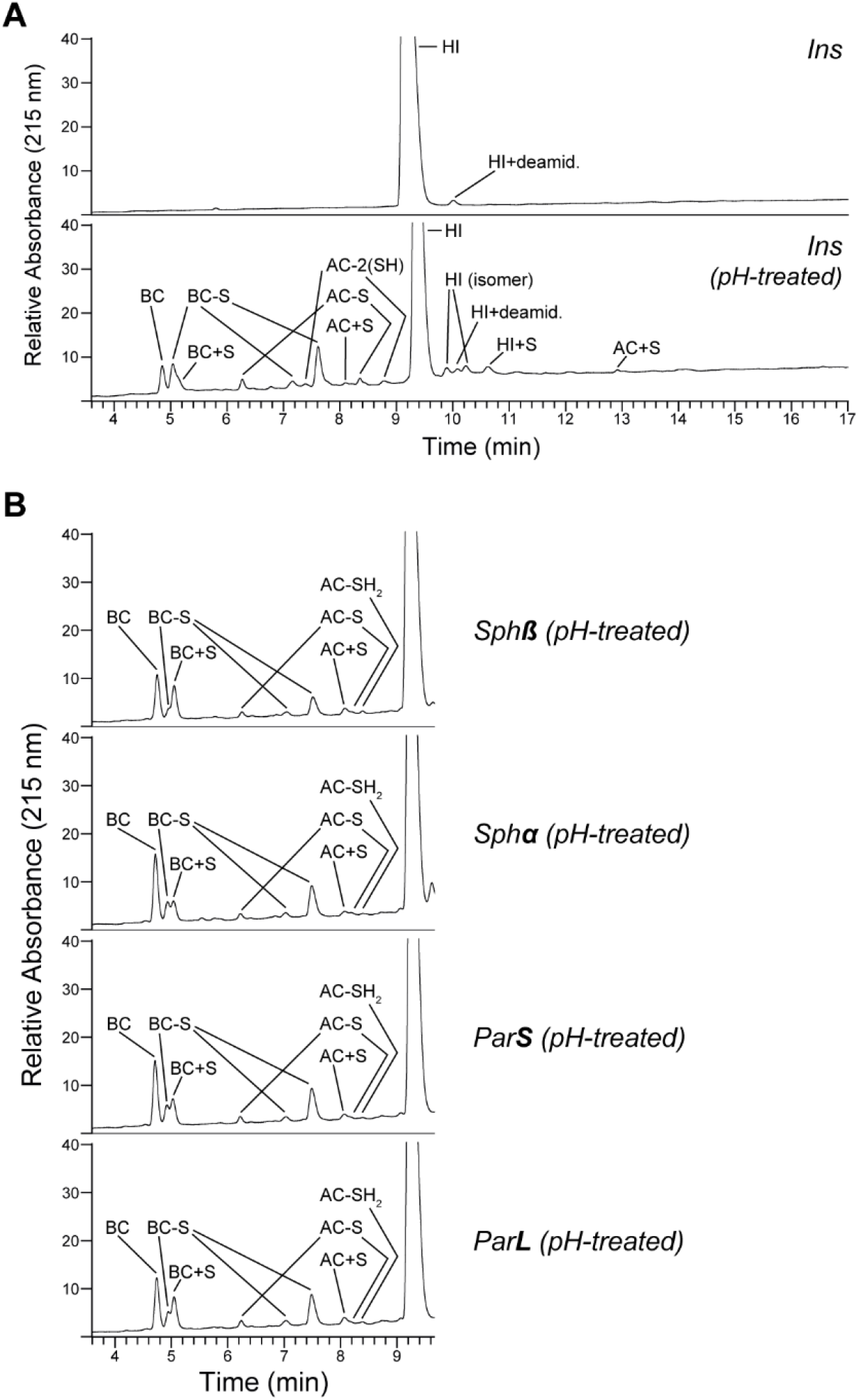
Disulfide bond breakage and scrambling from pH treatment. Representative LC-UV-MS chromatograms of the tested compounds showing disulfide bond breakage and scrambling when the high pH – low pH treatment used for aggregate dissociation is applied. UV detection at A215 nm are illustrated and the chemical modification products identified by mass spectrometry are annotated in the chromatograms for each sample. A unstressed human insulin samples, analysed either without (top) or with (bottom) the high pH – low pH treatement used for aggregate dissociation. B Products found in the dissociated aggregate samples eluting before the main peak of unmodified human insulin (HI).

**Figure S3.**
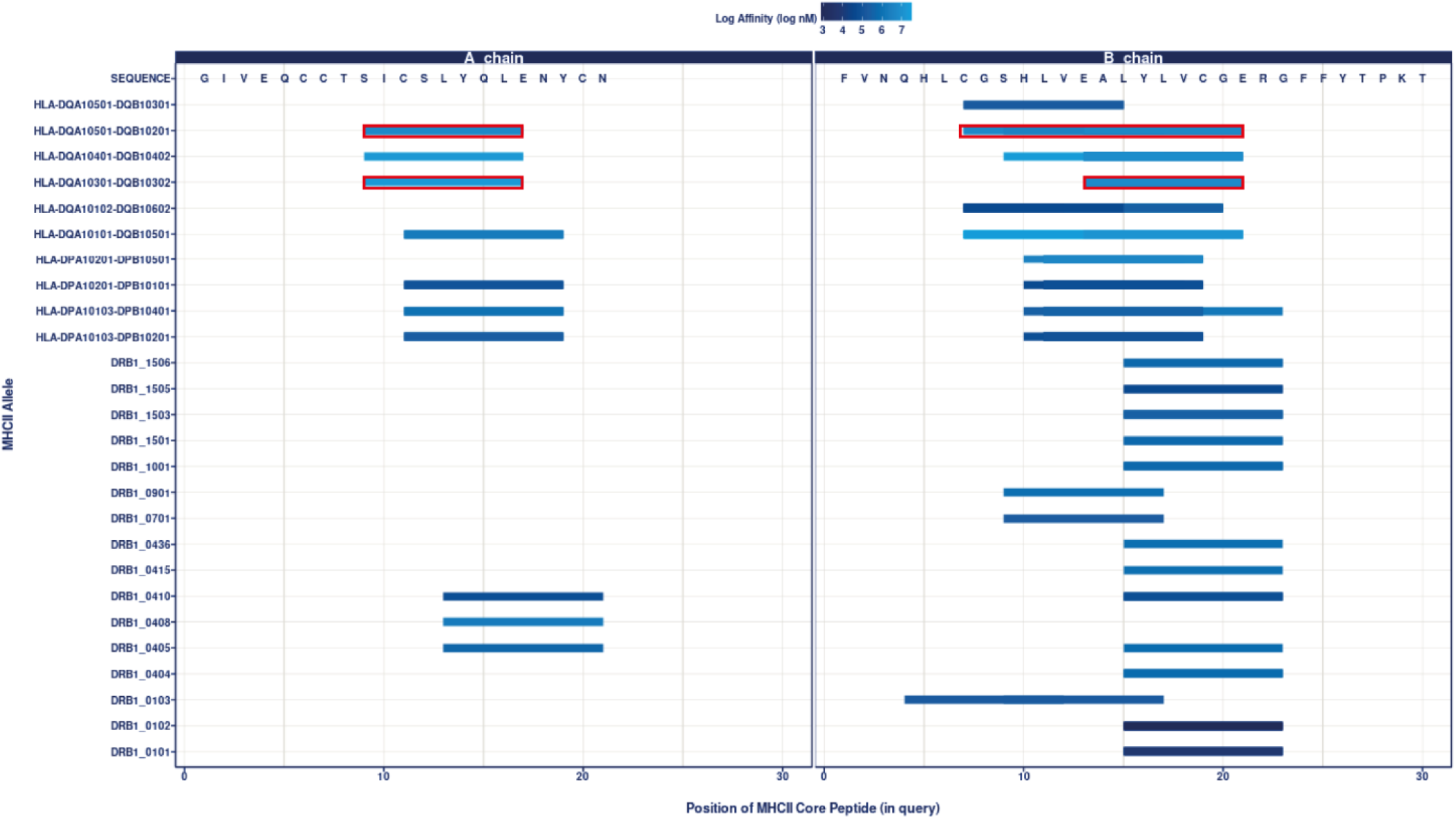
*In silico* analysis of HLA binding affinities. Binding affinities (log nM) of HLA in the A-chain or B-chain of human insulin predicting possible T-cell epitopes. Epitopes found for our chosen donor cohort are marked in red.

**Figure S4.**
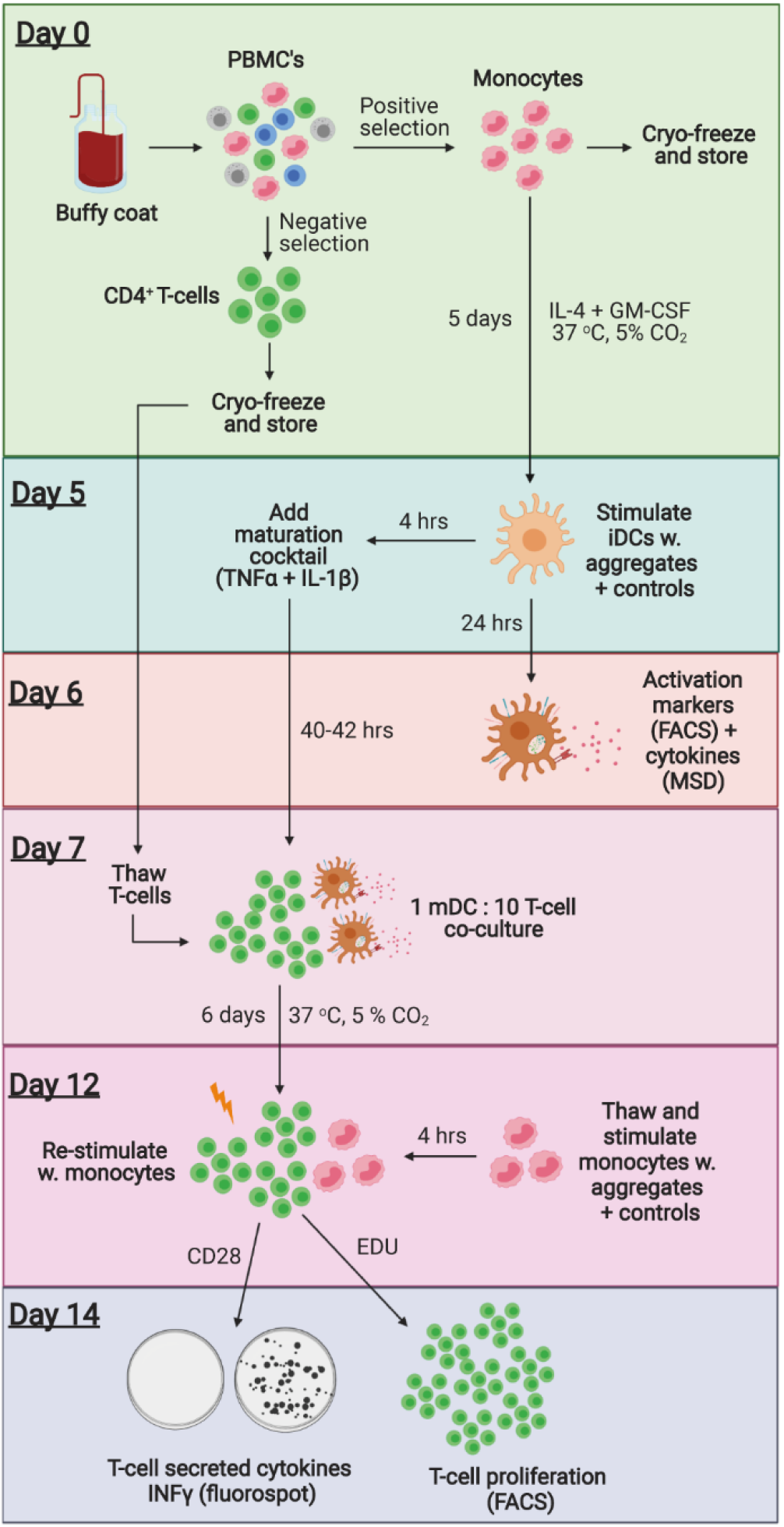
Graphical illustration of the *in vitro* immunogenicity assay protocol. Created with BioRender.com.

**Figure S5.**
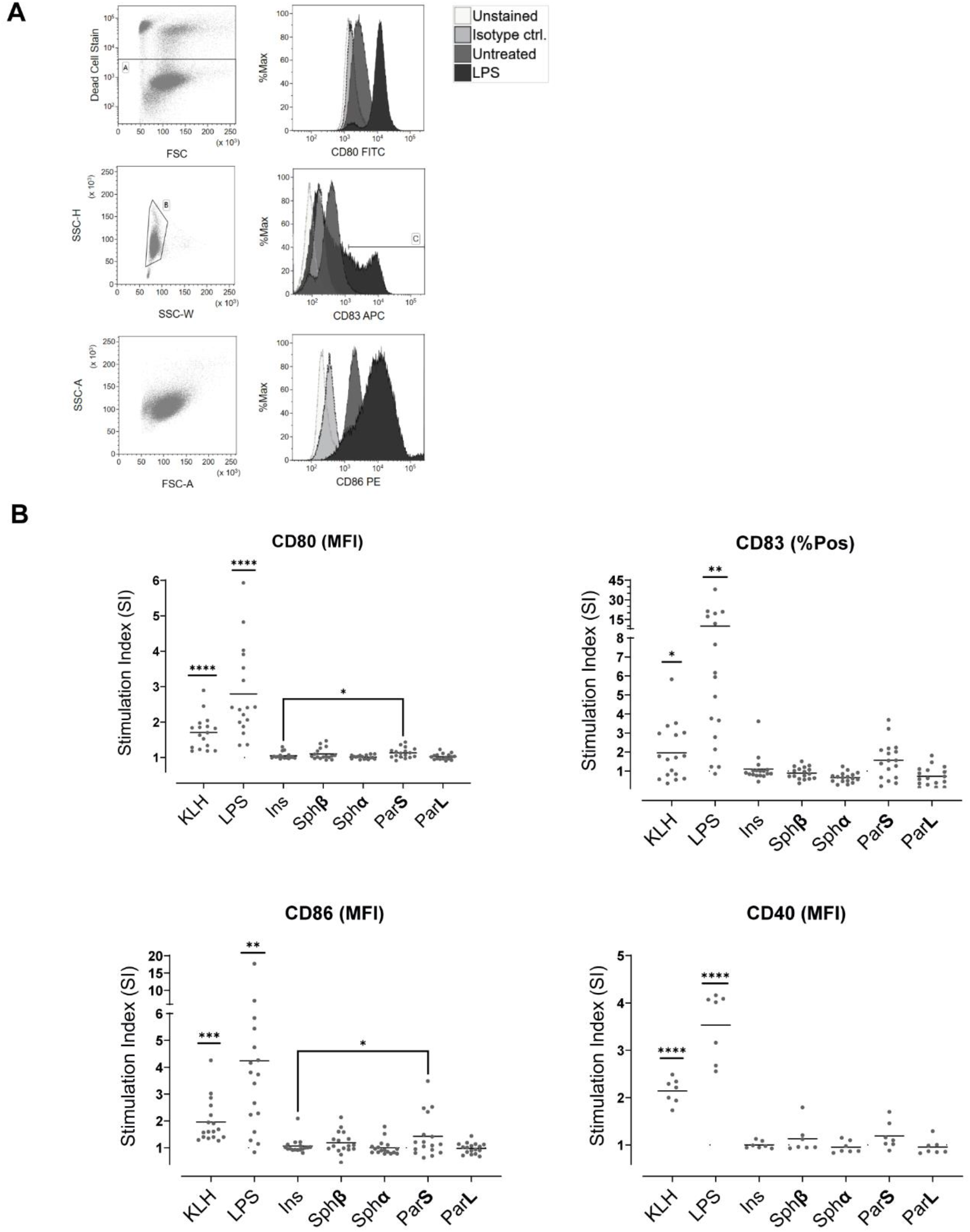
Flowcytometry analysis of monocyte-derived dendritic cell maturation. A Gating strategy for the data treatment, ensuring that data treatment was only performed on living, single cells. Representative histograms for maturation markers CD80, CD83 and CD86 shows that LPS (positive control) leads to an increase in fluorescence intensity compared to unstained, isotype control and untreated cells. B Data gathered for each maturation marker, where each data point represents a donor. For CD80, CD86 and CD40 the change in mean fluorescence intensity (MFI) was used, whereas percent positive cells (%Pos) was used for CD83. The stimulation index is calculated as reference to unstimulated cells, which is shown as the dotted line. Data information: There are N=17 donors included in the graphs for CD80, CD83 and CD86, and N=7 donors included for CD40.

**Figure S6.**
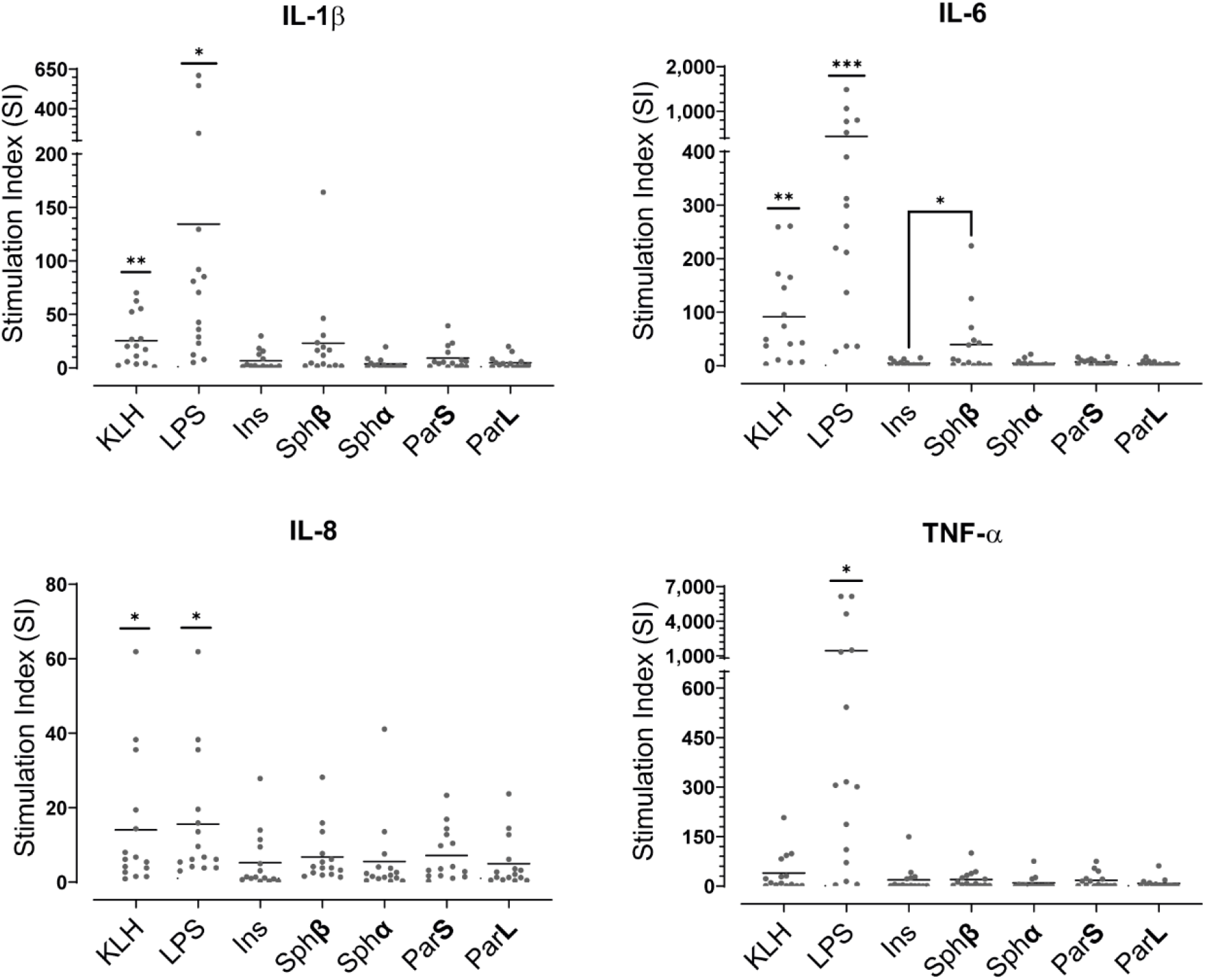
Multiplex cytokine analysis (IL-1β, IL-6, IL-8 and TNF-α) of the supernatant for monocyte-derived dendritic cell maturation using MSD. The stimulation index is calculated as reference to unstimulated cells, which is shown as the dotted line. Data information: There are N=15 donors included in the graphs.

**Figure S7.**
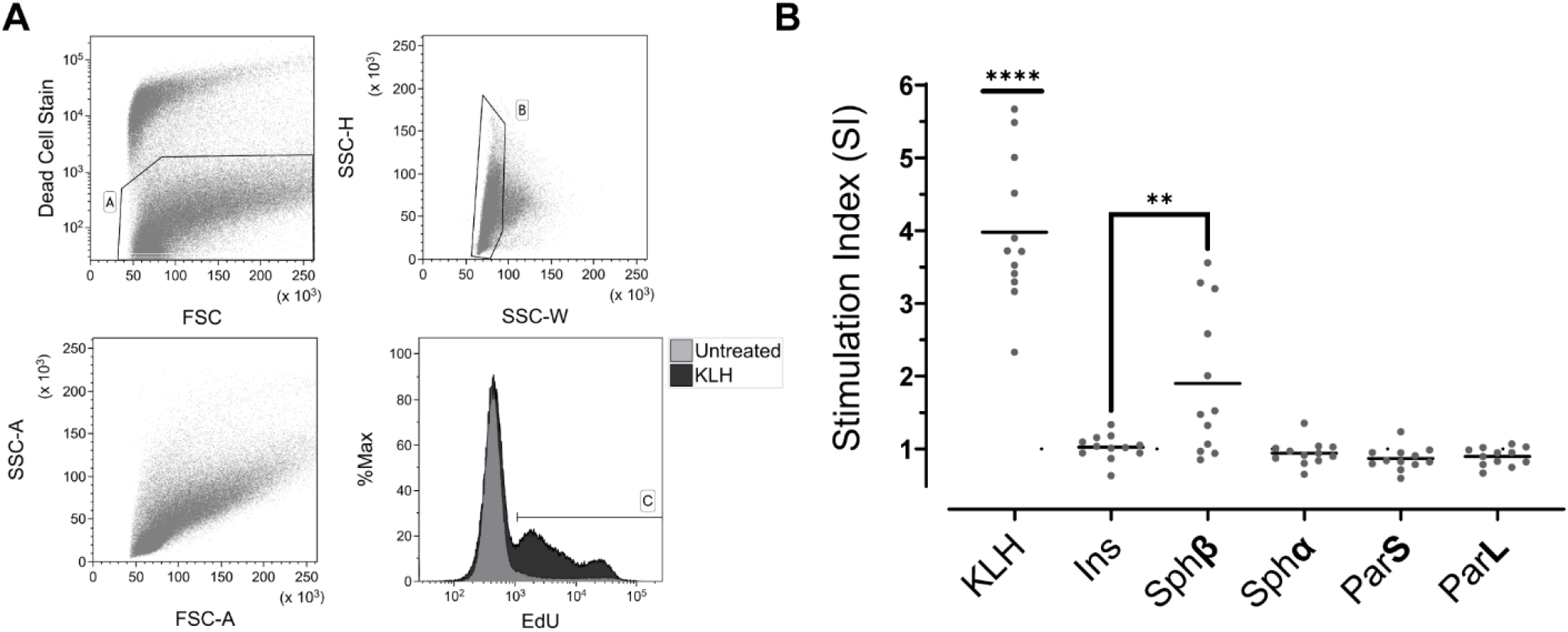
Flowcytometry analysis of CD4^+^ T-cell proliferation by EdU incorporation. A Gating strategy for the data treatment, ensuring that data treatment was only performed on living, single cells. Representative histogram for KLH (positive control) shows proliferation leads to an increase in cells with a higher fluorescence intensity compared to untreated cells (%Pos). B Data gathered, where each data point represents a donor. The stimulation index is calculated as reference to unstimulated cells, which is shown as the dotted line. Data information: There are N=12 donors included in the graph.

**Figure S8.**
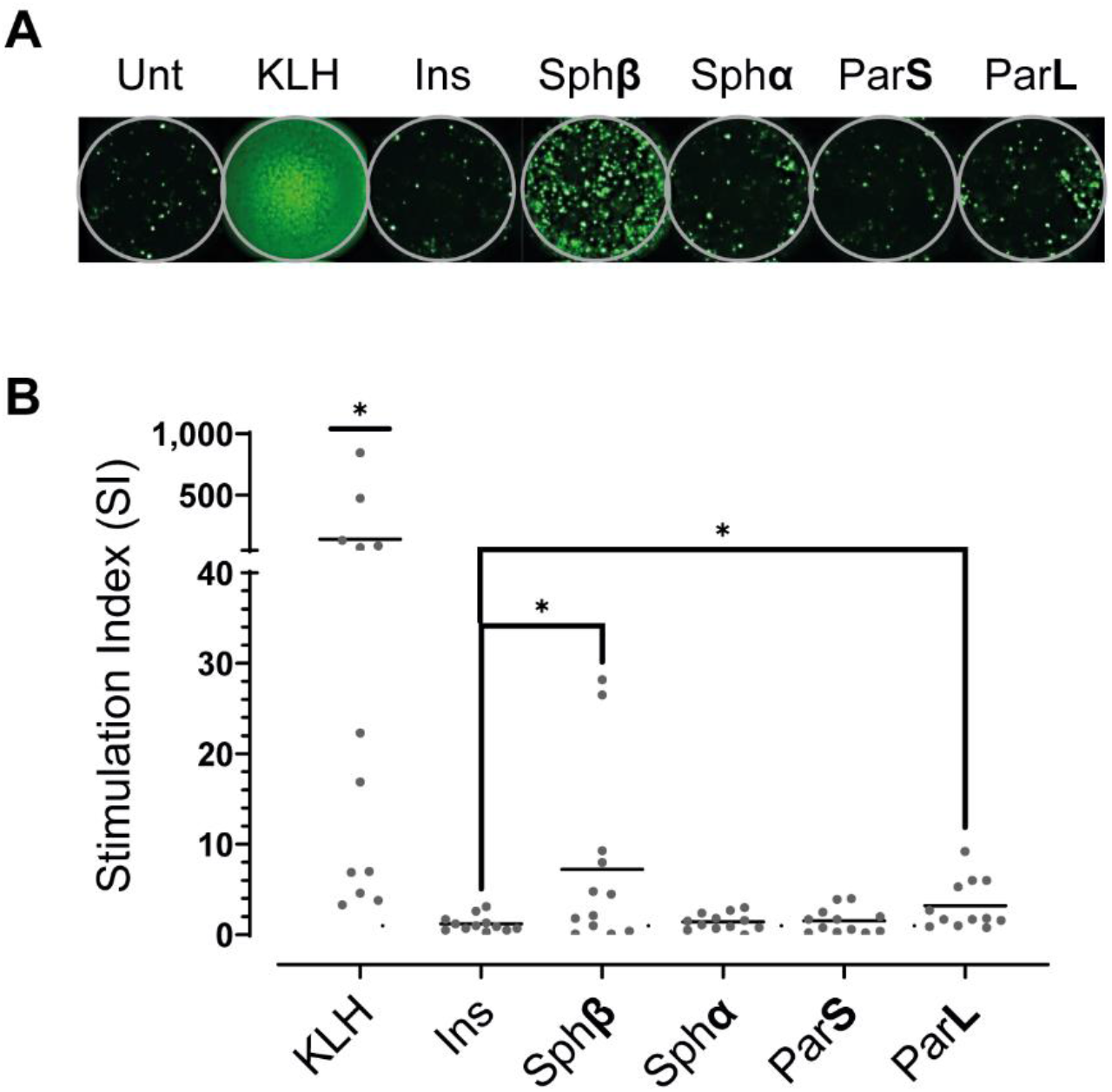
Fluorospot analysis of INF-γ secretion for CD4^+^ T-cell activation. A Fluorospot images from a representative donor showing the fluorescent signal for cells which has secreted INF-γ. The number of fluorescent cells are counted in each well and used for further analysis. B Data gathered, where each data point represents a donor. The stimulation index is calculated as reference to unstimulated cells, which is shown as the dotted line. Data information: There are N=12 donors included in the graph.

## Tables and their legends

**Table S1.** High resolution HLA II typing of study donor cohort. Shown in bold are alleles from donors whose HLA-DQ type has been associated with Type 1 diabetes.

| Donor ID | HLA-DQA1 |  | HLA-DQB1 |  |
| --- | --- | --- | --- | --- |
| 823 | **\*05:01** | **\*05:01** | **\*02:01** | *03:01 |
| 985 | *02:01 | **\*03:01** | **\*02:01** | **\*03:02** |
| 999 | **\*03:01** | **\*05:01** | *03:01 | *03:01 |
| 71 | *01:02 | **\*03:01** | **\*03:02** | *06:02 |
| 85 | *02:01 | **\*03:01** | **\*02:01** | *03:01 |
| 380 | *01:03 | **\*05:01** | *03:01 | *06:03 |
| 240 | *02:01 | **\*05:01** | **\*02:01** | **\*02:01** |
| 972 | *01:03 | **\*05:01** | **\*02:01** | *06:03 |
| 492 | *01:01 | **\*03:01** | **\*03:02** | *05:01 |
| 979 | **\*05:01** | **\*05:01** | **\*02:01** | **\*02:01** |
| 740 | *01:02 | **\*03:01** | **\*03:02** | *06:04 |
| 756 | *01:03 | **\*05:01** | *03:01 | *06:03 |
| 214 | **\*03:01** | **\*03:01** | **\*03:02** | **\*03:02** |
| 904 | *01:02 | *02:01 | **\*02:01** | *06:02 |
| 519 | **\*05:01** | **\*05:01** | **\*02:01** | **\*02:01** |
| 522 | **\*03:01** | **\*05:01** | **\*02:01** | **\*03:02** |
| 566 | *01:02 | **\*05:01** | **\*02:01** | *06:02 |

